# The mismatch in distributions of vertebrates and the plants that they disperse

**DOI:** 10.1101/120485

**Authors:** Jacob W. Dittel, Christopher M. Moore, Stephen B. Vander Wall

## Abstract

Little is known about how mutualistic interactions affect the distribution of species richness on broad geographic scales. It has been predicted that the richness of species involved in obligate mutualisms should be positively associated across their range. Whereas, if mutualisms are facilitative, the distribution of mutualists should be correlated with other factors. This study is the first study to compare the co-distribution of mutualist species in general and seed dispersal mutualisms specifically. We used geographic distributions of plant and animal mutualists to investigate the co-distribution and patterns of seed dispersal mutualisms. We found the mutualism between dispersers and plants does not account for the distribution of either group. In fact, there is a mismatch of richness between plants and the animals that disperse their seeds. Environmental factors are better predictors of both animal distribution and seed dispersal mutualisms across North America.

**Statement of authorship:** JD, CM, and SV conceived the original project idea. Plant data were compiled and analyzed by CM and SV, and JD compiled animal and environmental data. JD standardized and formatted all geographical data, and JD and CM performed all statistical analyses. JD wrote the first draft of the manuscript, and all authors contributed significantly to the revisions.

## INTRODUCTION

The broadest spatial scale of ecology examines patterns of diversity based on the geographic distributions of species (MacArthur 1972). Geographic distributions of species are important (i) to infer lower-level community-, population-, and physiological-based processes and (ii) as the ultimate comparison against which lower-level processes are compared (Bowyer *et al*. 1997; Somero 2005; Siefert *et al*. 2013). Indeed, there is a preponderance of studies of species richness at broad geographic scales (Oberdoff *et al*. 1995; Rahbek & Graves 2001; Hawkins *et al*. 2003a; Rahbek *et al*. 2007) that have facilitated our understanding of why species are found where they are, a central tenet within the domain of ecology (Scheiner & Willig 2008). Most commonly, species distributions are compared with environmental variables, which are presumed determinants of species distributions. Environmental variables are only one determinant of species' distributions, however; the other determinant—species interactions—is a key, understudied determinant of species’ distributions.

When species or guilds interact, we expect the geographic distributions of the pairs to be correlated. For pairwise interaction types where a species or guild benefits from another species or guild, we expect a positive relationship in distributions of abundance and richness due to the increase in fitness where they overlap (Svenning *et al*. 2014). This should be especially true in the case of mutualisms, where both sides of the interaction share an increase in fitness from being together (Bronstein 2015). In the case of seed dispersal mutualisms, for example, the distribution of plants and the animals that disperse their seeds should be similar. In terms of diversity, this further implies that the richness of plant species should maintain an equal or similar richness of animal species (Knops *et al*. 1999). Ultimately, this suggests for seed dispersal mutualisms that the richness of animal-dispersed plants *ought to be* correlated with the richness of their animal dispersers, and vice versa. To our knowledge, this prediction has never been tested on any appreciable geographic scale.

Seed dispersal is a critical stage in the life history of plants (Howe & Smallwood 1982; Herrera 1985; Terborgh 1990), and plants have evolved a variety of environmental and biotic methods to disperse their seeds. Of those plants that are actively dispersed by animal mutualists, the mutualistic interactions are presumed to be asymmetrical—plants are obligate mutualists dependent on guilds of dispersing animals, whereas animals are facultative mutualists partially depending on the plants they disperse. Many studies have described the richness of animal species (e.g., Jetz & Rahbek 2002; Kissling *et al*. 2007, 2009) and plant species (Kreft & Jetz 2007; Vander Wall & Moore 2016) at geographic scales and their relationship with various environmental variables. Yet despite the importance and commonality of these seed dispersal mutualisms, their distribution and prevalence are still poorly known globally (Vander Wall *et al*. submitted; Willson *et al*. 1990; Almeida-Neto *et al*. 2008). One factor limiting the testing of hypotheses of seed dispersal mutualism diversity, and the effects of plant diversity on animal diversity in general, has been the lack of plant data (Hawkins & Pausas 2004). Recently, Vander Wall & Moore (2016) described the distribution of plants dispersed by animal mutualists in North America. They found that plants with seed-dispersal mutualists appear to be more sensitive to environmental conditions than plants dispersed non-mutualistically but it is unknown whether the species richness of animal dispersers contributes to this pattern. In this study, we build upon Vander Wall and Moore’s (2106) finding to determine how the diversity of seed dispersing animals affects the diversity of animal-dispersed plants and/or if seed dispersal mutualism are related to environmental variables.

Given our theoretical predictions that species and guilds involved in mutualisms should be positively associated across their geographic distribution, we investigated the composite distribution of terrestrial bird and mammal seed dispersers in across North America. The ultimate goal of this study was to determine how the richness of seed dispersing animals varies across North America and how that composite distribution matches up with the distribution of the plants that they disperse. To do so, we investigated whether the richness of seed dispersing animals is correlated with broad ecological and environmental variables, including the richness of the plants that they disperse. We hypothesize that if the mutualisms are obligate, as is presumed between plants being obligate mutualists of the seed dispersers, then the richness distribution of the plants would show a stronger relationship with seed dispersers compared with environmental variables. Alternatively, if the mutualisms between plants and seed dispersing animals is facultative or diffuse, as is presumed between the animals that disperse the plants, then we expect a stronger relationship between the richness distribution of animals and environmental variables compared with the plants. We methodologically used GIS and spatial statistics to investigate if plant mutualist abundance or environmental factors were correlated with seed dispersing animals and seed dispersing mutualisms. We surprisingly found that there is a mismatch between the richness of seed dispersing animals and the plants they disperse. Environmental variables were more often correlated with the distribution of seed dispersing animals and better explained the distribution of seed dispersing mutualisms than the richness of either plant or animal mutualists.

## MATERIALS AND METHODS

### Assignment of species to seed dispersing guilds

To determine how the distribution of vertebrate seed dispersers compares to the plants that they disperse, we first assigned the birds and mammals of North America (north of Mexico) to two seed dispersing guilds: frugivorous and scatter-hoarding seed dispersers. Animals were considered frugivorous seed dispersers if they consume fruits containing seeds as a significant portion of their diet, and the seeds remain viable after being either regurgitated or passed through the digestive tract. Scatter-hoarding of seeds, which frequently results in a mutualism with plants, is limited to the bird family Corvidae and the mammal order Rodentia in North America (Vander Wall 1990). A species was considered a scatter-hoarder if seeds are a significant portion of its diet, it scatter-hoards them in soil, and there is a reasonable expectation that some of those seeds germinate. Hereafter, we use the terms frugivore and scatter-hoarder to mean species that are mutualist seed dispersers. Full details of species assignment can be found in appendix S1a.

### Data acquisition and preparation

We prepared for our analyses by first creating comparable datasets. We had four groups of data: animal mutualists, environmental variables, plant mutualists, and the difference between animal and plant mutualists. The animal mutualists consisted of seven subguilds: all animal mutualists, frugivorous, scatter-hoarding, frugivorous mammals, frugivorous birds, scatter-hoarding rodents, and scatter-hoarding birds. Species data consisted of a polygon of the geographic distribution involved in the type of mutualism. Bird distribution data were obtained from BirdLife International and NatureServe (Ridgley *et al*. 2007; BirdLife International and NaturServe 2014). Mammal distribution data were obtained from the International Union for Conservation of Nature Red List (IUCN 2012). These polygons were overlaid, richness was summed, and the resulting file was rasterized to generate animal mutualist species richness at each grid cell of a master raster.

The environmental variables consisted of four datasets: mean actual evapotranspiration (mm/yr; hereafter AET), elevation (m), mean precipitation (mm/year; hereafter precipitation) and latitude (degrees). AET was obtained from the Global-AET Database (Trabucco & Zomer 2010), elevation was obtained from Natural Earth (2017), and precipitation was obtained from Bioclim (Hijmans *et al*. 2005). These environmental variables were chosen because they have been found to be important predictors in previous studies of species distributions (Pearson & Dawson 2003). AET is a proxy for terrestrial productivity (Mackey & Currie 2001), and has been found to be associated with bird and plant distributions (Karr 1976; Hawkins *et al*. 2003b; Kissling *et al*. 2009). Precipitation is predicted to be important for scatter-hoarding behaviors, with the behavior being more frequent in semi-arid and arid ecosystems rather than mesic ones (Vander Wall & Jenkins 2011). Lastly, we included elevation because there are large elevational gradients in western North America and species richness generally decreases with an increase in elevation (Rahbek 1995). Each environmental variable was obtained as a raster and bilinear interpolation was used to conform the extent and resolution of the original raster to our master raster.

The plant mutualists consisted of seven datasets: plant mutualists, plants in frugivory mutualism, plants in scatter-hoarding mutualism, plants in frugivory mutualism with mammals, plants in frugivory mutualism with birds, plants in scatter-hoarding mutualism with rodents, and plants in scatter-hoarding mutualism with birds. Each plant dataset was obtained from a previous study that identified plants in dispersal mutualism at 197 sites across North America, north of Mexico (Vander Wall & Moore 2016). For this study, we used the plant species richness per site that are dispersed by frugivores or scatter hoarders. (See Vander Wall & Moore (2016) for details on their methods and dispersal mode determination.) We interpreted the point values and estimated values across our geographic range of interest. Specifically, we used ordinary kriging to interpolate values to our master raster using R library, automap (Hiemstra *et al*. 2009). The difference between animal and plant mutualists were for the same seven modes listed above. Because the means and variances were very different between animal and plant mutualists, we calculated z-scores (eqn. 1) between them, and subtracted the z-score of plant from animal mutualists at each point to create a value (Zdiff) used in data analysis described below.

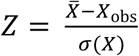
Where 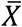 is the mean richness, X_obs_ is the richness at the specific point, and σ(*X*) is the standard deviation of the richness of either animal or plant mutualists. Further explanation on how the rasters for the Z_diff_ were created, including details of extent can be found in Appendix S1b.

### Data analysis

Data analysis was broken into three major categories: all seed dispersing mutualists, frugivorous animals, and scatter-hoarding animals. Within each major category, we had two general comparisons: (1) animal mutualists and plant mutualists and (2) Z_diff_ and environmental variables. Due to high heteroscedasticity in the plant richness, data-weighted least squares regression models were created using the squared residuals of area adjusted plant richness as weights. Despite richness being count data, a Poisson distribution was not necessary because the data did not deviate from a normal distribution. Spatial autocorrelation is a common occurrence in range map and atlas survey data (Dormann *et al*. 2007) and was present in the environmental variables, but none of the richness of either plants or animals. To adjust for spatial autocorrelation, generalized least squares models (GLS) were built for each comparison using a Gaussian spatial correlation. Data were transformed with a natural log when necessary.

We then conducted two types of Monte Carlo simulations to test the hypothesis that there is no relationship between data within our groups of variables: a complete randomization and a spatially-structured randomization. The complete randomization permuted grid cells across the continent, which allowed us to test the hypothesis that observations are random. The structured randomization statistically fit a spatial autocorrelation model (variogram), and then generated a random field with the same degree of spatial autocorrelation using the Random Fields package in R (Schlather *et al*.2013, 2015). Given that we know that geographic data are spatially autocorrelated, this allowed us to test the hypothesis that, given our observed levels of spatial autocorrelation, the observed data are random. We calculated Spearman’s correlation coefficient, *p*, for each of 1000 interactions for the complete and structured randomizations. We then calculated the proportion of the complete or structured randomizations that were more extreme than the observed correlation, *ρ*^*^ as an estimated *p*-value, 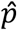. R code used to conduct the analyses can be found at the following url: https://github.com/dispersing/SpatialRandomizations. Every complete randomization failed to detect a random distribution; thus, rendering the analysis uninformative. (Supplementary Information S6–S9 shows the full results of the complete randomizations.) Therefore, every Monte Carlo simulation reference henceforth is specific to the spatially-structured tests.

Lastly, because so little is known about the factors contributing to seed-dispersal mutualism distributions we performed classifications and regression trees (CART) after our initial analysis to better understand the structure of the data and identify factors that warrant future investigation. For this data exploration, we used R library, rpart (Therneau *et al*. 2015) using the ANOVA method. All analysis was performed in program R (R Development Core Team 2017).

## RESULTS

### Distribution of seed dispersing animals

We identified 183 animal species in North America that have a seed dispersing mutualism with plants either via frugivory or scatter-hoarding. Seed dispersing animals were most speciose in the southwestern portions of North America from the southern portion of the Colorado Plateau desert region and further north, east of the Rocky Mountains, to the southern Rocky Mountain-prairie border (Fig. 1). There was no relationship between the richness of all mutualist animals and the plants they disperse; instead richness of all animal mutualists decreases with an increase in latitude (*F*_1,195_ = 207, *p* < 0.001, Fig. 2). Results from the regression models and the Monte Carlo simulations can be found in Table 1. The primary split in the CART model for all mutualists was at ∼50°N latitude. At latitudes ≥ 50°N, the richness of animal mutualists is correlated with plant richness (*F*_1,15_ = 25.27*,p* < 0.001), but not at latitudes < 50°N (*F*_1,177_ = 2.47, *p* = 0.12).

**Figure 1:**
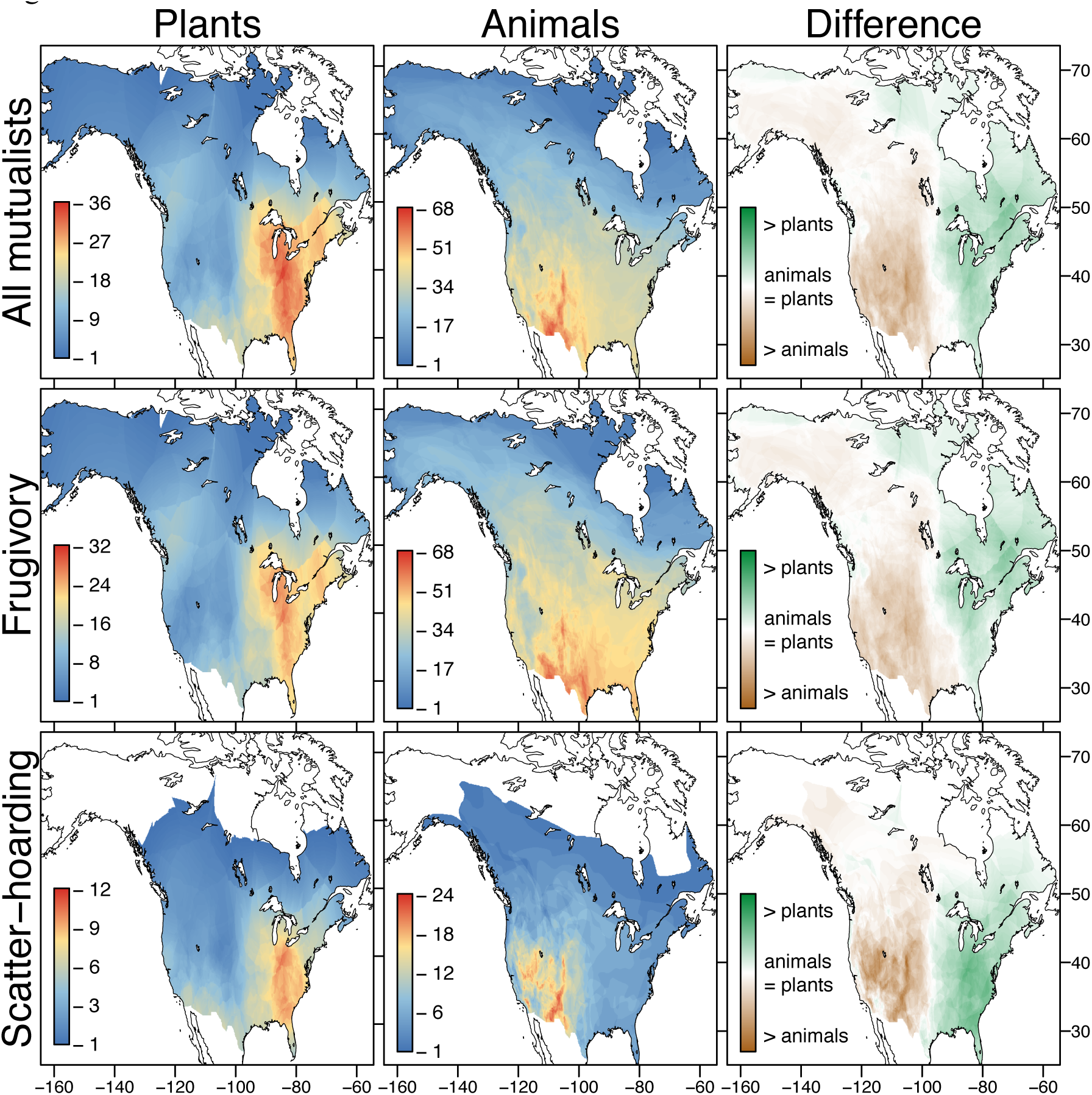
Distribution of seed dispersal mutualism richness. The left panels are the distribution of plants involved in each guild of seed dispersal mutualism after (Vander Wall *et al*. submitted). The middle panel is the richness distribution of animals involved in each guild of seed dispersal mutualisms, and the right panel is the difference between plant and animal richness (Z_diff_) involved in guild of mutualisms. In the left and middle panels, the legend represents the number of plants or animals at any given locations. While in the right panel the legend represents the relative difference in richness between plants and animals.

**Figure 2:**
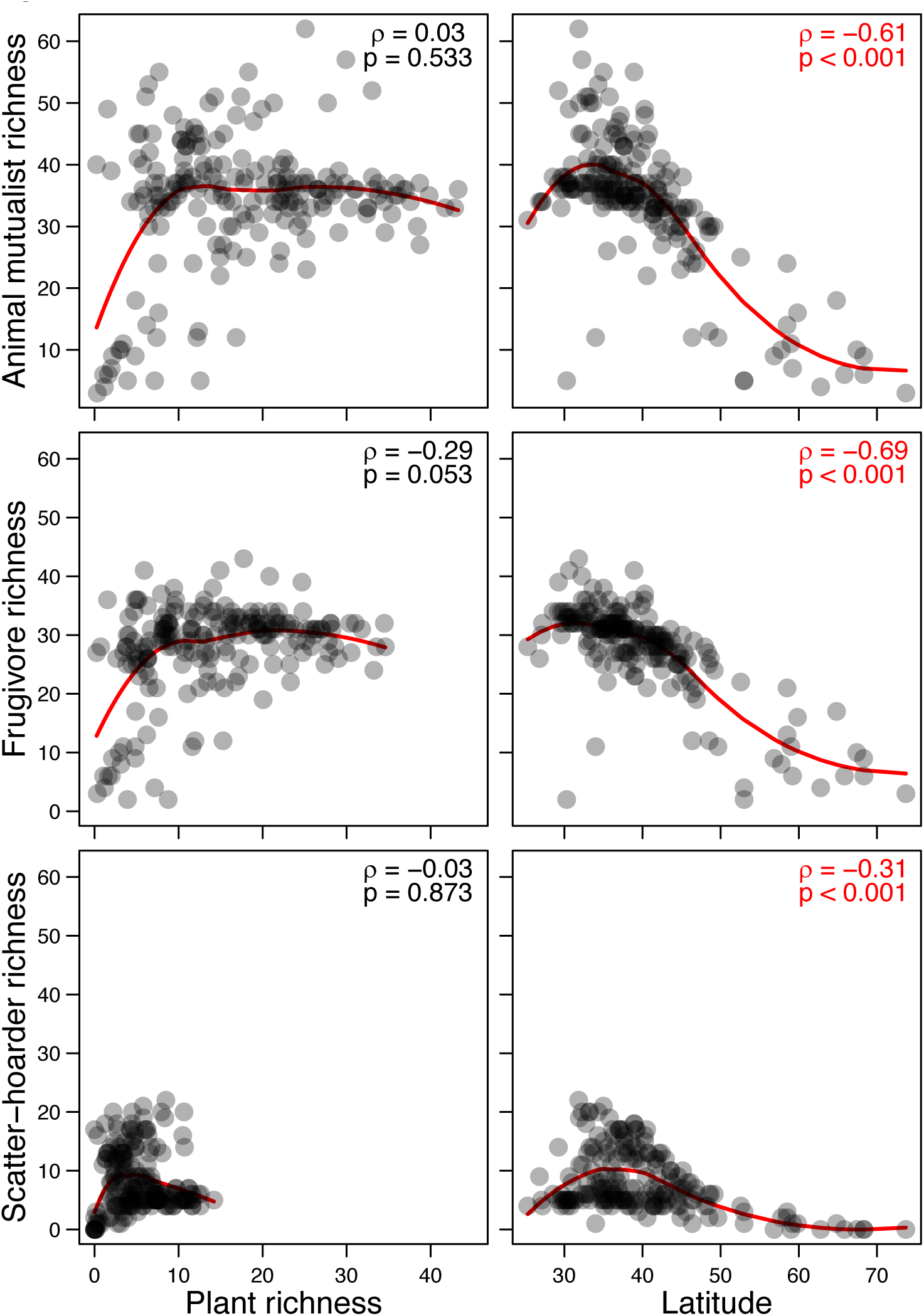
The richness of animals involved in seed dispersing mutualisms as a function of the plants they disperse (left column) and latitude (right column). Each plot has the Spearman correlation coefficient (*ρ*^*^) andp-value of the linear model; red lettering signifies a significant linear model. The only significant interactions were between the richness of all mutualists and frugivores with latitude (all mutualists: *F*_1,195_= 207, *p* <0.001, frugivores: *F*_1,195_ = 262.6, *p* < 0.001). The red line is the LOWESS smoothing curve to reveal the internal structure of the distribution of data points.

**Table 1:** Statistical results of regression models and Monte Carlo simulations for subguild richness (dependent variable) and the richness of the plants they disperse (independent variable). For the regression models, the table includes the *F*-statistic, *p*-value, effect size (*β*), and *R^2^* for each model. Monte Carlo simulations include the predicted *p*-value (
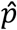
) and Spearman’s correlation coefficient (*ρ*\*).

| Dependent variable | Independent variable | Regression models |  |  |  | Monte Carlo simulations |  |
| --- | --- | --- | --- | --- | --- | --- | --- |
| | | *F*-statistic | *p*-value | Effect size ($\beta$) | $R^2$ | $\hat{p}$ | $\rho^*$ |
| All mutualists | All plant mutualists | 0.39 | 0.534 | 0.33 | 0.00 | 0.02 | 0.66 |
| All frugivores | Plants dispersed by frugivores | 3.79 | 0.053 | 0.71 | 0.01 | 0.01 | 0.67 |
| Frugivorous birds | Plants dispersed by frugivorous birds | 1.05 | 0.306 | 0.37 | 0.00 | 0.01 | 0.62 |
| Frigivorous mammals | Plants dispersed by frugivorous mammals | 1.24 | 0.327 | 0.24 | 0.00 | 0.04 | 0.44 |
| All scatter-hoarders | Plants dispersed by scatter-hoarders | 0.03 | 0.873 | -0.02 | -0.01 | 0.03 | 0.59 |
| Scatter-hoarding birds | Plants dispersed by scatter-hoarding birds | 0.33 | 0.566 | -0.04 | 0.00 | 0.03 | 0.47 |
| Scatter-hoarding rodents | Plants dispersed by scatter-hoarding rodents | 0.34 | 0.561 | 0.05 | 0.00 | 0.04 | 0.58 |

A total of 88 animal species were determined to participate in a frugivorous seed-dispersal mutualism, 65 species of birds and 23 species of mammals (Table S2). Frugivore richness is highest in the southwestern portions of North America, specifically in the Colorado Plateau semi-desert region, east to the Southwest plateau and dry steppe region (Fig. 1). Richness is relatively low in the Great Basin and Mojave deserts, with the lowest richness in in the far north tundra; richness of frugivorous animals decreases with increasing latitude (*F*_1,195_ = 262.6, *p* < 0.001, Fig. 2). There is no relationship with frugivorous animal richness and the richness of the plants they disperse (Table 1). CART models again showed a primary split at ∼50°N latitude, and when the data was divided at that point, there were similar relationships between richness as we found for all mutualists. At latitudes ≥ 50°N, frugivore richness and the richness of the plants they dispersed are correlated (*F*_1,15_ = 18.88*,p* < 0.001). However, at latitudes < 50°N, this relationship disappears (*F*_1,177_ = 1.71, *p* = 0.19). Frugivorous bird and frugivorous mammal richness are similarly correlated negatively with latitude but not correlated with plant richness (Table 1).

Lastly, we identified a total of 102 animal species as scatter-hoarders involved in seed dispersal mutualisms; 10 species of birds and 92 species of rodents (Table S3). As with frugivorous animals, scatter-hoarder richness is concentrated in the southwestern North America. Scatter-hoarder richness is highest in Chihuahuan desert region, with richness hotspots in the Great Basin and Mojave deserts. The Sonoran Desert has a surprisingly low scatter-hoarder richness (Fig. 1). Richness is lower in eastern North America with the lowest regions being in the Adirondacks and northern tundra. Richness decreases with an increase in latitude (*F*_1,195_ = 39.01, *p* < 0.001), but is not correlated with plant richness (Table 1, Fig. 2). CART models do not show a ∼50°N latitude split (in fact, latitude is not a primary split in the data at all), instead the primary split occurs at ∼900 m elevation. Further exploratory analysis of the data did not provide any relationships between scatter-hoarder richness and the plants they disperse between the high- and low-elevation groups. Scatter-hoarding rodents are similarly negatively correlated with latitude and are not correlated with plant richness. Conversely, scatter-hoarding birds are not correlated with plant richness nor latitude (Table 1)

### Seed dispersal mutualisms

There was a clear mismatch of richness between seed dispersers and the plants that they disperse (Fig. 1) with the highest richness of plants dispersed by animals being in eastern North America, while the highest richness of animal dispersers being in western North America. The divide is approximately 100°W longitude for both guilds of seed dispersers, and the two guilds combined. Indeed, in all CART models, longitude is the second split in the data further suggesting its importance.

There was no relationship between the Z_diff_ of all mutualists and precipitation, AET, nor latitude (Table 2) However, there was a negative relationship with median elevation (Fig. 3) and the Z_dif_ of all mutualists. As elevation increases, there was a larger proportion of animal richness comparted to plant richness. Monte Carlo simulations supported the observed relationship with elevation was different from random, and supported our findings of no relationships between other variables (Table 2). The sub-panels in Figure 3 show the results of the Monte Carlo simulations which were largely consistent with our regression models.

**Table 2:** Statistical results of GLS models and Monte Caro simulations for mutualisms (Z_dif_; dependent variable) and environmental variables (independent variables). For spatial regression models, the table includes whether there was a spatial correlation in the data, how it was corrected for, *F*-statistic,*p*-value, effect size (*β*), and *R^2^* for each model. Monte Carlo simulations include the predicted *p*-value (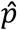) and Spearman’s correlation coefficient (*ρ*\*). A dagger (†) signifies that the effect size is in units per 100 mm/yr AET and a double dagger (††) signifies that the effect size is in units per 100 m elevation.

| Independent variable | Dependent variable | Spatial regression models |  |  |  |  | Monte Carlo simulations |  |
| --- | --- | --- | --- | --- | --- | --- | --- | --- |
| | | Spatial correlation | $F$-statistic | $p$-value | Effect size ($\beta$) | $R^2$ | $\hat{p}$ | $\rho^*$ |
| AET | All mutualists | Gaussian | 3.57 | 0.06 | 0.08<sup>†</sup> | 0.03 | 0.07 | 0.39 |
|  | All frugivores | Gaussian | 2.07 | 0.15 | 0.06<sup>†</sup> | -0.02 | 0.18 | 0.24 |
|  | All scatter-hoarders | Gaussian | 247.53 | 0.00 | 0.34<sup>†</sup> | 0.23 | 0.00 | 0.68 |
| Elevation | All mutualists | Gaussian | 160.42 | 0.00 | -0.09<sup>††</sup> | 0.08 | 0.00 | -0.55 |
|  | All frugivores | Gaussian | 84.45 | 0.00 | -0.07<sup>††</sup> | -0.02 | 0.00 | -0.49 |
|  | All scatter-hoarders | Gaussian | 224.85 | 0.00 | -0.18<sup>††</sup> | 0.19 | 0.00 | -0.54 |
| Latitude | All mutualists | Gaussian | 1.63 | 0.20 | 0.02 | 0.07 | 0.34 | 0.15 |
|  | All frugivores | Gaussian | 12.41 | 0.00 | 0.04 | 0.20 | 0.22 | 0.19 |
|  | All scatter-hoarders | Gaussian | 0.62 | 0.43 | 0.01 | -0.01 | 0.29 | -0.14 |
| Precipitation | All mutualists | Gaussian | 0.32 | 0.57 | 0.11 | 0.01 | 0.13 | 0.25 |
|  | All frugivores | Gaussian | 46.33 | 0.00 | 0.70 | 0.02 | 0.22 | 0.19 |
|  | All scatter-hoarders | Gaussian | 109.06 | 0.00 | 1.19 | 0.00 | 0.04 | 0.36 |

**Figure 3:**
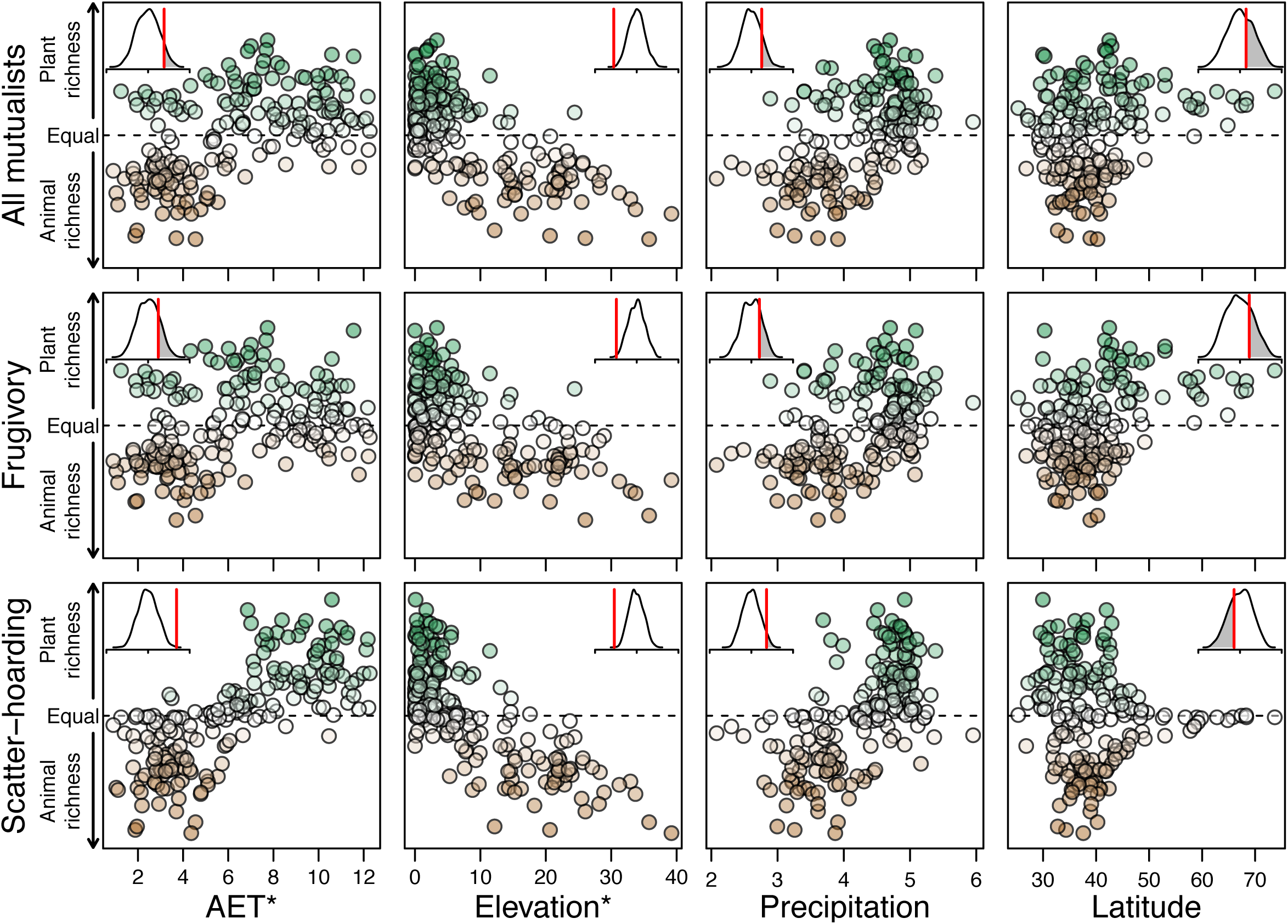
The relationships between environmental variables (rows) and differences in the richness of plant and animal mutualists (Z_diff_; columns). All statistics and relationships can be found in Table 2. The scatterplots show the environmental variable on the x axis and the difference in plant and animal mutualist richness of on the *y* axis. From left to right, the environmental variables represent AET (mm/yr), elevation (m), precipitation (log(mm/yr)), and latitude. An asterisk (*) next to the variable implies that the values are in the hundreds. Each scatterplot is colored according to greater plant richness (green), equal plant and animal richness (white), or greater animal richness (brown). We further included subplots of the spatially-structured randomizations to show how extreme the observed correlation (Spearman’s *ρ*; vertical red line) was compared with 1,000 randomizations (density plot). The subplots have ticks for values for correlation values *ρ*^*^ = -1, 0, and 1.

There was no relationship between Z_diff_ of frugivores and precipitation, AET, or latitude; but there was with median elevation (Table 2, Fig. 3). Similarly, there were no relationships between Z_diff_ of frugivorous birds and precipitation, AET, latitude, median elevation; nor between Z_diff_ of frugivorous mammals and precipitation, AET, latitude, nor median elevation (Table S4). Monte Carlo simulations largely supported our findings again (Fig. 3, sub-panels) but suggested that our data was different from random for elevation (*ρ* = -0.49,*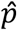*= 0), suggesting that we observed fewer frugivorous animals at higher elevations than plants dispersed by them.

The Z_diff_ of scatter-hoarding animals was also not correlated with latitude (Table 2). However, there were relationships with AET, precipitation, and median elevation. As AET or precipitation increases there are proportionately more plants dispersed by scatter-hoarders than scatter-hoarders (Fig. 3). The proportion of scatter-hoarders increased compared to the plants they disperse with an increase in median elevation (Fig. 3). The Z_di_ff of scatter-hoarding birds follows the same pattern as the whole guild. There was no relationship between the Z_diff_ of scatter-hoarding birds and latitude, but there were relationships with precipitation, AET, and median elevation (Table S4). There was an increase in the proportion of scatter-hoarders as AET decreased or elevation increased. The Z^diff^ of scatter-hoarding rodents also was correlated with precipitation, AET, median elevation (Table S4). The proportion of scatter-hoarding rodents increased with a decrease in precipitation, a decrease in AET, or an increase in elevations. The Z_diff_ of scatter-hoarding rodents was not correlated with latitude. Monte Carlo simulations again largely corroborated our findings, with the only contradicting result involving precipitation. Simulations suggest there was a relationship with precipitation for all scatter-hoarders, with more scatter-hoarders being found in areas of less precipitation (*ρ* = -0.04,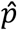=0.04, Fig. 3 sub-panels).

## DISCUSSION

There was an apparent mismatch between the richness of seed dispersing animals and the plants that they disperse across North America. We hypothesized that because of the increased fitness between two species or guilds of mutualists that we would expect positive relationships of species richness between both sides of the mutualism, depending on the degree of co-dependency (i.e., facultative-to-obligate). Although we found positive relationships between all subguilds of seed dispersing animals and seed-dispersed plants, the richness of both plants and animals were better explained by environmental variables than by the richness of each other. This result was surprising and suggests that diversity of mutualistic partners do not necessarily beget diversity of one another.

Species richness of frugivores, scatter-hoarders, and all seed dispersing animals combined decreased with an increase in latitude (Fig. 2). This pattern matches the pattern observed for the plants that they disperse (Vander Wall *et al*. submitted; Vander Wall & Moore 2016). The increase of species richness with decreasing latitude was not a surprising result as the generality of the latitudinal diversity gradient has been found to be robust (Hillebrand & Thomas 2004). However, when hoarding birds are considered alone, they did not show a significant latitudinal gradient (*F*_1,194_ = 3.08, *p* = 0.08). This result was likely because the data are limited to one small, generalist family of birds (Corvidae) that have their center of diversity in the north temperate zone.

As with previous studies (Karr 1976; Hawkins *et al*. 2003b; Kissling *et al*. 2009), animal species richness was correlated with environmental variables (data not shown). Surprisingly, the difference in relative abundance of animals and plants in seed dispersal mutualisms was not strongly correlated with environmental variables. One exception was that the proportion of plants dispersed by scatter-hoarding animals increased with AET (Fig. 5), a variable often found important for predicting species richness at large scales. Seed disperser mutualisms appear to be more frequent (i.e., more plants dependent on scatter-hoarders) in more productive environments. Our Monte Carlo simulations also suggested that scatter-hoarding mutualism should be correlated with precipitation (which is correlated with AET) and we found that richness of scatter-hoarding rodents was correlated with precipitation. These results are congruent with the predictions made by Vander Wall & Jenkins (2011) on why western species of chipmunks *(Tamias)* have adapted scatter-hoarding in arid and semi-arid western North America, but not in the mesic eastern North America. However, the proportion of scatter-hoarders was positively correlated with median elevation (Fig. 3) and does not follow the global pattern; a decrease in richness with an increase in elevation (Rahbek 1995). Richness is believed to decrease at higher elevations partially due to a decrease in productivity, smaller land area, and harsher climates leading to higher extinction and lower dispersal due to greater distances between suitable habitat (Rahbek 1995; Rowe 2009; Wu *et al*. 2013). The relationship with elevation may be due to the fact that in southwestern North America, where the majority of seed-disperser richness was found, net productivity actually increases with elevation into the montane forests before decreasing again above tree line (Whittaker & Niering 1975). Since plants are positively correlated with AET, this may influence the relationship with elevation.

The distribution of animal species richness contrasts with the richness distribution of the plant species being dispersed by those animals (Figs. 3, 4). Plants dispersed by frugivores and scatter-hoarding animals have their highest richness around the Great Lake regions and southeastern parts of North America in general. This mismatch of species richness between plants and the animals that disperse them is clearly seen in Fig. 1 with animal richness being greater in southwestern North America and plant richness being greater in southeastern North America. A possible explanation for this enigma is that seed dispersing animals might be far more numerous in the southeastern United States, where there are relatively few species, compared to the southwestern part of the continent where abundance may be lower, but species richness is higher.

The strength of coevolution between individual plant and vertebrate disperser species has been suggested to be diffuse (Thompson 1982; Wheelwright & Orians 1982; Herrera 1985). In particular seed dispersal communities of plant and animal species have found that species interactions are often asymmetrical, variable in time and space, and non-obligate (Janzen 1980; Wheelwright 1988; Bascompte & Jordano 2007). This is particularly driven by generalists animal species, which interact with multiple plant species causing high complementarity and trait convergence (Guimaraes Jr. *et al*., 2011). Most, if not all, of the animal species in North America that we considered seed disperser mutualists would fall under this definition of generalists as they have wide diet breadths. The plants dispersed by animals in North America are also generalists as fruits and seeds have evolved to attract a variety of dispersers and not any one species in particular. Diffuse interactions inhibit strong directional coevolution and lead to the diffuse patterns we witness.

The current distribution of plants and animals in North America has changed over the last 18,000 years (Ray & Adams 2001). The last glacial maximum dramatically altered species distributions across North America, and it may be that there has not been enough time since the last glacial maximum for coevolutionary selection pressures to form or be strong enough to be detected at coarse spatial scales. It is also probable that animals have migrated faster than the plants they disperse because of the diffuse relationship between animals and plants, in addition to the short period since the last glacial maximum. Davis *et al*. (1986) and Woods & Davis (1989) have shown that some animal-dispersed plants have not reached their potential distributions since the last glacial maximum, despite their dispersers being common across the plant’s potential geographic range.

Two major limitations of the study were (i) the assumption that abundance is uniform across a species range and (ii) a mismatch of scale between occurrence and environmental data. First, assumptions of studies using species range data are that the abundance of a species is uniform across its range, that all species have equal abundances, and that abundances are high enough throughout the range for the species to be an effective part of the community. This latter point, in this case, means that each species is an effective disperser of plants wherever it occurs. These assumptions are rarely met (Hurlbert & Jetz 2007), but occurrence maps are typically the only data on species occurrence available at large spatial scales, and if maps are constructed in a similar way, they can provide insights into the richness of species in a region (Rocchini *et al*. 2011).

Secondly, Hurlbert & Jetz (2007) also suggested that a mismatch of scale between occurrence data and environmental variables can lead to erroneous results. Instances of mismatch often occur when species occurrence data (generally course resolution) is overlaid onto climatic variables (generally finer resolution). We believe the concerns of mismatch are minimal for this study as the overarching aim was to identify the distribution of animals in comparison to the plants that they disperse. Analyses with climatic variables were chosen based on previous findings and hypotheses and the data were taken at the coarsest scale available to match occurrence data as best as possible. As with similar studies, the purpose of these analyses is to identify broad patterns of distribution with the goal of providing focal points for finer scale studies and not to suggest detailed patterns.

This study is the first to compare the collective distribution of animals involved in seed dispersal mutualisms to the distribution of the plants they disperse. The distribution of plants nor animals accounts for the distribution of either group. In fact, there is an apparent mismatch of richness between plants and the animals that disperse their seeds (Figs. 1, 2). As with animal-dispersed plants (Vander Wall & Moore 2016) environmental variables, particularly latitude, better describes the richness distribution of animal mutualists. In the case of seed dispersal mutualisms, median elevation was correlated with all mutualists and scatter-hoarding, additionally, scatter-hoarding was correlated with AET suggesting that environmental factors, such as productivity may play a key role in the distribution of the mutualism. Further work is sorely needed to better understand the effect of climate on distributions of seed dispersing animals. However, with this data of seed dispersing animal distributions, we can now identify locations that warrant further study either to understand better seed-dispersal mutualisms or the factors that influence the distribution of the plants and animals involved in these mutualisms. The biggest challenge to further understanding many of these observed patterns has been the lack of appropriate data (Hawkins & Pausas 2004), and we hope this study will serve as a stepping stone to further discoveries.

## Acknowledgements

We’d like to thank M. Matocq, C. Feldman, W. Longland, and S. Mensing for providing comments on a previous draft of the manuscript. We’d also like to thank Bioclim, Global-AET, Natural Earth, Bird Life International, NatureServe, IUCN and the for the data used in this study

## Appendix S1

### a)Full description of species assignment to seed-dispersing guilds

To determine how the distribution of vertebrate seed dispersers compares to the plants that they disperse, we first assigned the birds and mammals of North America (north of Mexico) to two seed-dispersing guilds: frugivorous and scatter-hoarding seed dispersers. Animals were considered frugivorous seed dispersers if they consume fruits containing seeds as a significant portion of their diet, and the seeds remain viable after being either regurgitated or passed through the digestive tract. We excluded animals that eat lots of fleshy fruit but that do not disperse seeds, like the northern cardinals *(Cardinalis cardinalis)* and ruffed grouse *(Bonasa umbellus)*, which eat fruit, but possess gizzards that destroy seeds by grinding them up. Additionally, animals that disperse seeds but do so infrequently (e.g. most members of the family Vireonidae), were excluded, as they are unlikely to influence plant-disperser coevolution in a significant way. Scatter-hoarding of seeds, which frequently results in a mutualism with plants, is limited to the bird family Corvidae and the mammal order Rodentia in North America (Vander Wall 1990). A species was considered a scatter-hoarder if seeds are a significant portion of its diet, it scatter-hoards them in soil, and there is a reasonable expectation that some of those seeds germinate.

Animals were identified to be frugivorous or scatter-hoarders through a combination of personal knowledge and the literature. For species that we suspected of being seed dispersers, we consulted the literature. This included Bird species accounts (American Ornithologist Union), Bent’s Life History of Birds series (Bent, 1919-1968), and Mammalian species accounts (American Society of Mammalogists). In some cases we surveyed the primary literature. If no source provided evidence for frugivory or scatter-hoarding of seeds, we excluded the species from the list. The taxonomy of all species is based on current species lists published for birds and mammals of North America (American Ornithologists’ Union 1998; Wilson & Reeder 2005). Our list of seed-dispersing vertebrates is intended to be comprehensive but the inclusion or exclusion of certain species is debatable. But we maintain that the inclusion or exclusion of a few species is unlikely to change our results in any meaningful way.

### b)Raster Creation

All of the rasters were then masked to our geographic range of interest (North America, north of Mexico) using major land areas (largely excluding islands) from CIA World DataBank II (https://www.evl.uic.edu/pape/data/WDB/) in R library, mapdata (Brownrigg 2016). The master raster has a resolution of 1221 × 1077 longitude by latitude (573933 terrestrial cells evaluated), and an extent of 164.4167 ° W–54.41667°W and 25.06667°N – 73.56667°N. Data were rasterized, overlaid, and projected using Program R (R Core Team 2014) with R libraries, maptools (Lewin-Koh *et al*. 2011), raster (Hijmans 2014), sp (Pebesma & Bivand 2005; Bivand *et al*. 2008), and rgdal (Keitt *et al*. 2014). Composite distribution maps were made for all mutualists (frugivores + scatter-hoarders), all frugivorous animals, frugivorous birds, frugivorous mammals, all scatter-hoarding animals, scatter-hoarding birds, and scatter-hoarding mammals.

For initial data analysis, we extracted the animal mutualist richness from the 197 sites used in Vander Wall & Moore (2016). To obtain richness, we used the extract function from the R library, Raster (Hijmans 2014). This function works by extracting specified data from a raster at given points. The latitude and longitude of the center (mean latitude and longitude) of each site was used as x, y coordinates for data extraction. The center of each site was deemed acceptable because the range maps of the included animals are not detailed enough to show significant differences within a given study site. Furthermore, we reviewed the species list for both frugivorous and scatter-hoarding animals at each site to eliminate species that were supposedly present according to their range maps but were unlikely actually present due to elevational or ecological limitations.

**Table S2:**
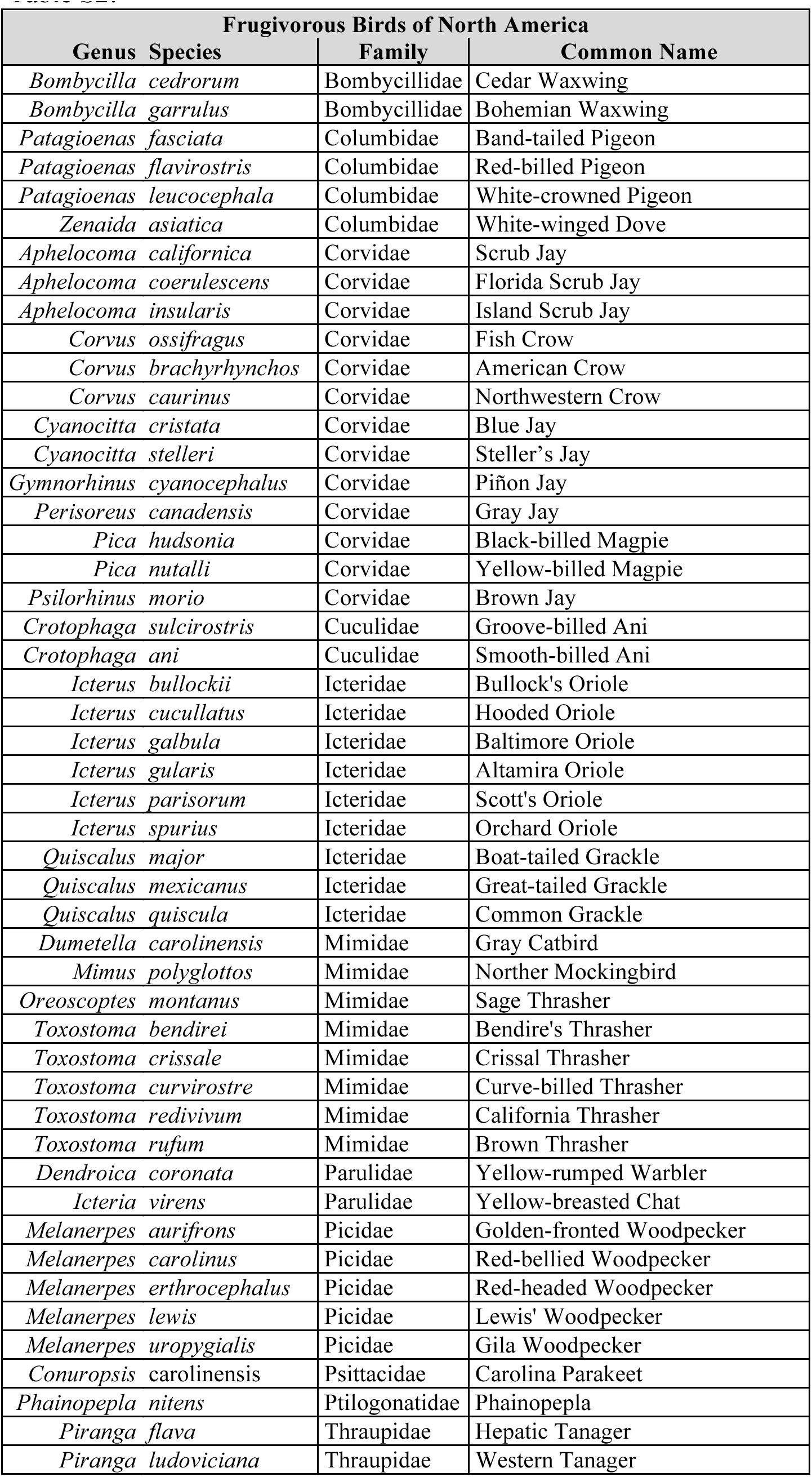
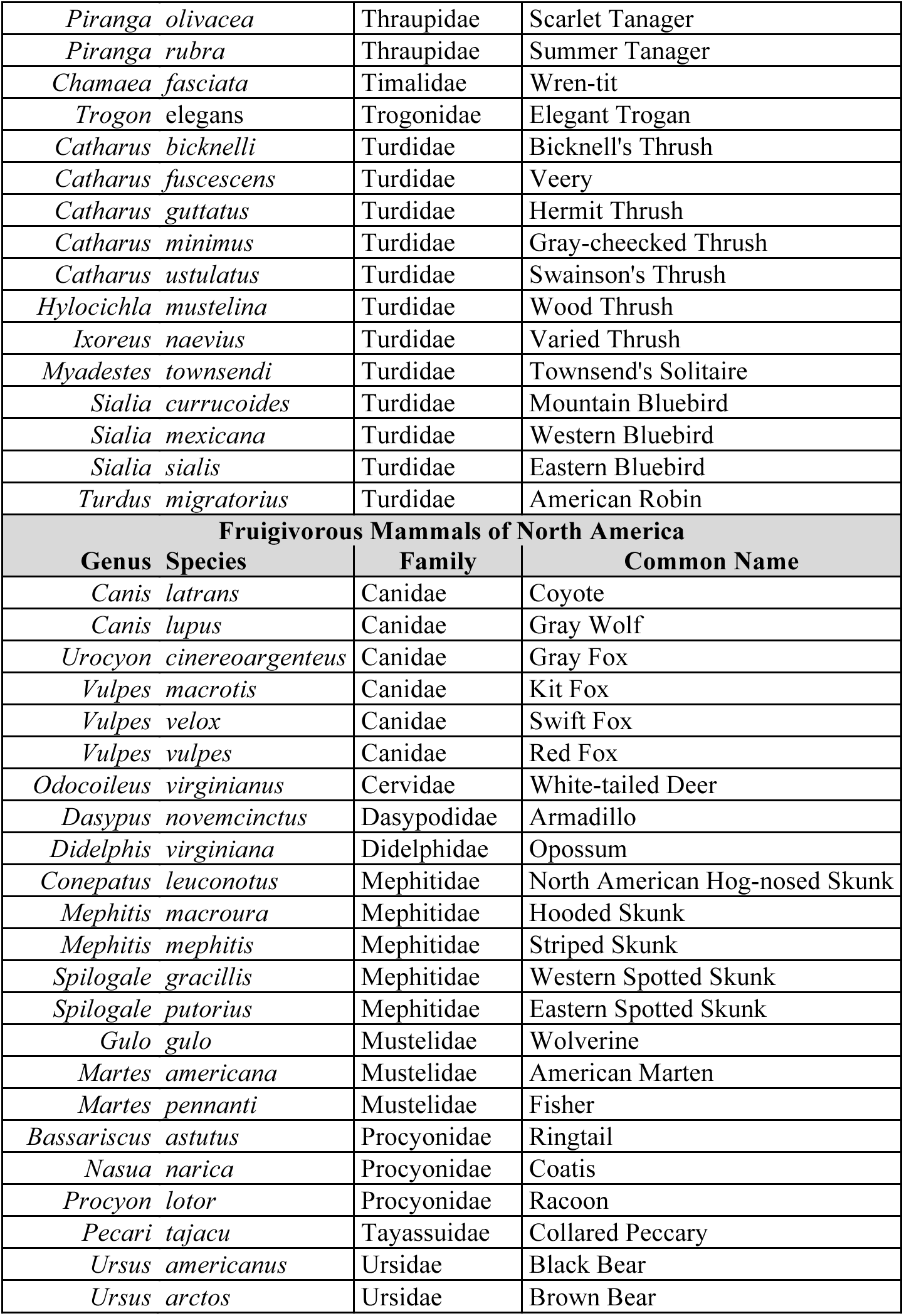
List of frugivorous animals broken up into frugivorous birds (top) and frugivorous mammals (bottom). Species are listed alphabetically by family then genus, order of species does not insinuate a species ability to disperse seeds via frugivory.

| Frugivorous Birds of North America |  |  |  |
| --- | --- | --- | --- |
| Genus | Species | Family | Common Name |
| *Bombycilla* | *cedrorum* | Bombycillidae | Cedar Waxwing |
| *Bombycilla* | *garrulus* | Bombycillidae | Bohemian Waxwing |
| *Patagioenas* | *fasciata* | Columbidae | Band-tailed Pigeon |
| *Patagioenas* | *flavirostris* | Columbidae | Red-billed Pigeon |
| *Patagioenas* | *leucocephala* | Columbidae | White-crowned Pigeon |
| *Zenaida* | *asiatica* | Columbidae | White-winged Dove |
| *Aphelocoma* | *californica* | Corvidae | Scrub Jay |
| *Aphelocoma* | *coerulescens* | Corvidae | Florida Scrub Jay |
| *Aphelocoma* | *insularis* | Corvidae | Island Scrub Jay |
| *Corvus* | *ossifragus* | Corvidae | Fish Crow |
| *Corvus* | *brachyrhynchus* | Corvidae | American Crow |
| *Corvus* | *caurinus* | Corvidae | Northwestern Crow |
| *Cyanocitta* | *cristata* | Corvidae | Blue Jay |
| *Cyanocitta* | *stelleri* | Corvidae | Steller's Jay |
| *Gymnorhinus* | *cyanocephalus* | Corvidae | Piñon Jay |
| *Perisoreus* | *canadensis* | Corvidae | Gray Jay |
| *Pica* | *hudsonia* | Corvidae | Black-billed Magpie |
| *Pica* | *nutalli* | Corvidae | Yellow-billed Magpie |
| *Psilorhinus* | *morio* | Corvidae | Brown Jay |
| *Crotophaga* | *sulcirostris* | Cuculidae | Groove-billed Ani |
| *Crotophaga* | *ani* | Cuculidae | Smooth-billed Ani |
| *Icterus* | *bullockii* | Icteridae | Bullock's Oriole |
| *Icterus* | *cucullatus* | Icteridae | Hooded Oriole |
| *Icterus* | *galbula* | Icteridae | Baltimore Oriole |
| *Icterus* | *gularis* | Icteridae | Altamira Oriole |
| *Icterus* | *parisorum* | Icteridae | Scott's Oriole |
| *Icterus* | *spurius* | Icteridae | Orchard Oriole |
| *Quiscalus* | *major* | Icteridae | Boat-tailed Grackle |
| *Quiscalus* | *mexicanus* | Icteridae | Great-tailed Grackle |
| *Quiscalus* | *quiscula* | Icteridae | Common Grackle |
| *Dumetella* | *carolinensis* | Mimidae | Gray Catbird |
| *Mimus* | *polyglottos* | Mimidae | Norther Mockingbird |
| *Oreoscoptes* | *montanus* | Mimidae | Sage Thrasher |
| *Toxostoma* | *bendirei* | Mimidae | Bendire's Thrasher |
| *Toxostoma* | *crissale* | Mimidae | Crissal Thrasher |
| *Toxostoma* | *curvirostre* | Mimidae | Curve-billed Thrasher |
| *Toxostoma* | *redivivum* | Mimidae | California Thrasher |
| *Toxostoma* | *rufum* | Mimidae | Brown Thrasher |
| *Dendroica* | *coronata* | Parulidae | Yellow-rumped Warbler |
| *Icteria* | *virens* | Parulidae | Yellow-breasted Chat |
| *Melanerpes* | *aurifrons* | Picidae | Golden-fronted Woodpecker |
| *Melanerpes* | *carolinus* | Picidae | Red-bellied Woodpecker |
| *Melanerpes* | *erthrocephalus* | Picidae | Red-headed Woodpecker |
| *Melanerpes* | *lewis* | Picidae | Lewis' Woodpecker |
| *Melanerpes* | *uropygialis* | Picidae | Gila Woodpecker |
| *Conuropsis* | *carolinensis* | Psittacidae | Carolina Parakeet |
| *Phainopepla* | *nitens* | Ptilonotidae | Phainopepla |
| *Piranga* | *flava* | Thraupidae | Hepatic Tanager |
| *Piranga* | *ludoviciana* | Thraupidae | Western Tanager |

|  |  |  |
| --- | --- | --- |
| *Piranga olivacea* | Thraupidae | Scarlet Tanager |
| *Piranga rubra* | Thraupidae | Summer Tanager |
| *Chamaea fasciata* | Timalidae | Wren-tit |
| *Trogon elegans* | Trogonidae | Elegant Trogon |
| *Catharus bicknelli* | Turdidae | Bicknell's Thrush |
| *Catharus fuscescens* | Turdidae | Veery |
| *Catharus guttatus* | Turdidae | Hermit Thrush |
| *Catharus minimus* | Turdidae | Gray-cheeked Thrush |
| *Catharus ustulatus* | Turdidae | Swainson's Thrush |
| *Hylocichla mustelina* | Turdidae | Wood Thrush |
| *Ixoreus naevius* | Turdidae | Varied Thrush |
| *Myadestes townsendi* | Turdidae | Townsend's Solitaire |
| *Sialia currucoides* | Turdidae | Mountain Bluebird |
| *Sialia mexicana* | Turdidae | Western Bluebird |
| *Sialia sialis* | Turdidae | Eastern Bluebird |
| *Turdus migratorius* | Turdidae | American Robin |
| **Fruigivorous Mammals of North America** |  |  |
| **Genus Species** | **Family** | **Common Name** |
| *Canis latrans* | Canidae | Coyote |
| *Canis lupus* | Canidae | Gray Wolf |
| *Urocyon cinereoargenteus* | Canidae | Gray Fox |
| *Vulpes macrotis* | Canidae | Kit Fox |
| *Vulpes velox* | Canidae | Swift Fox |
| *Vulpes vulpes* | Canidae | Red Fox |
| *Odocoileus virginianus* | Cervidae | White-tailed Deer |
| *Dasypus novemcinctus* | Dasypodidae | Armadillo |
| *Didelphis virginiana* | Didelphidae | Opossum |
| *Conepatus leuconotus* | Mephitidae | North American Hog-nosed Skunk |
| *Mephitis macroura* | Mephitidae | Hooded Skunk |
| *Mephitis mephitis* | Mephitidae | Striped Skunk |
| *Spilogale gracillis* | Mephitidae | Western Spotted Skunk |
| *Spilogale putorius* | Mephitidae | Eastern Spotted Skunk |
| *Gulo gulo* | Mustelidae | Wolverine |
| *Martes americana* | Mustelidae | American Marten |
| *Martes pennanti* | Mustelidae | Fisher |
| *Bassariscus astutus* | Procyonidae | Ringtail |
| *Nasua narica* | Procyonidae | Coatis |
| *Procyon lotor* | Procyonidae | Raccoon |
| *Pecari tajacu* | Tayassuidae | Collared Peccary |
| *Ursus americanus* | Ursidae | Black Bear |
| *Ursus arctos* | Ursidae | Brown Bear |

**Table S3:**
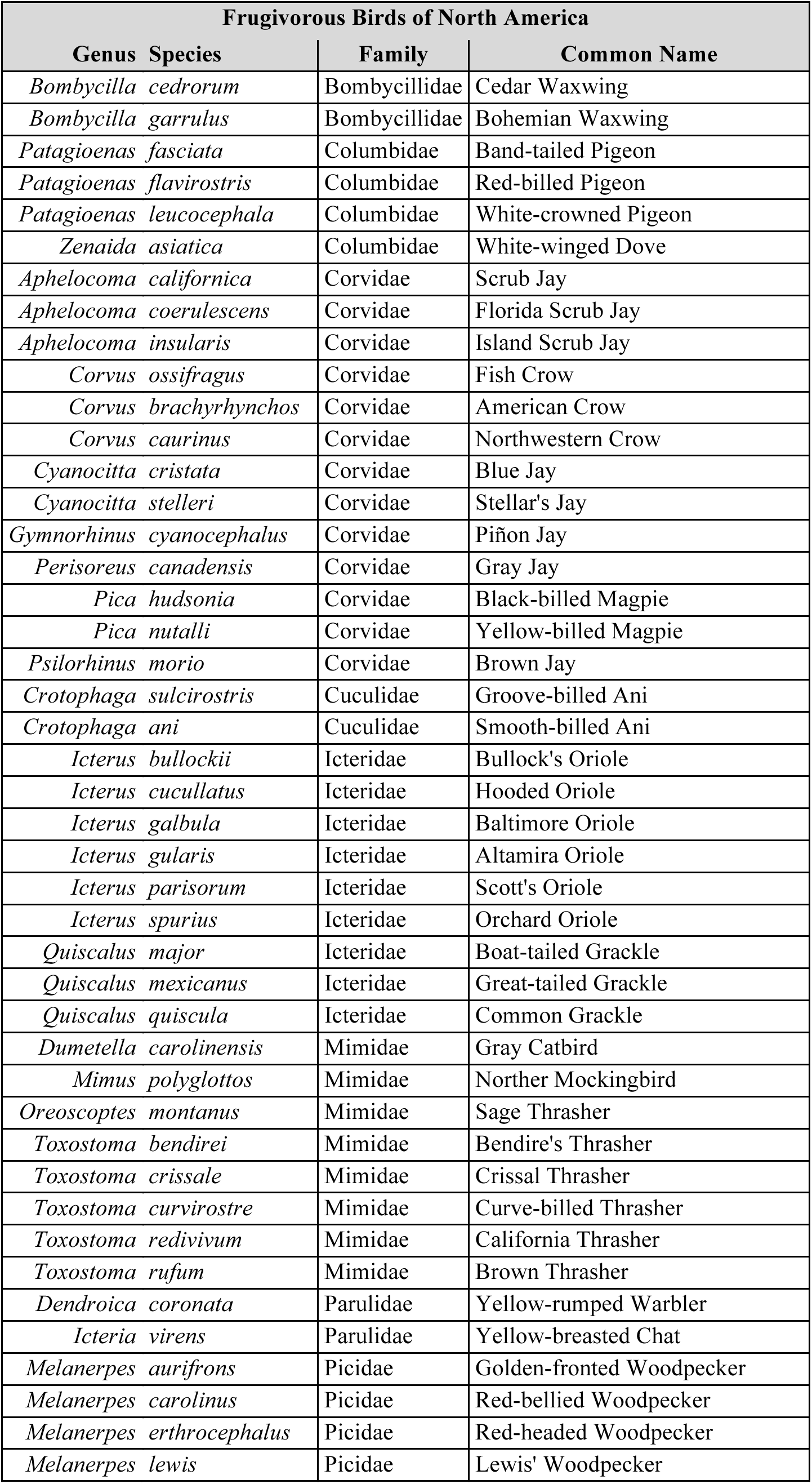
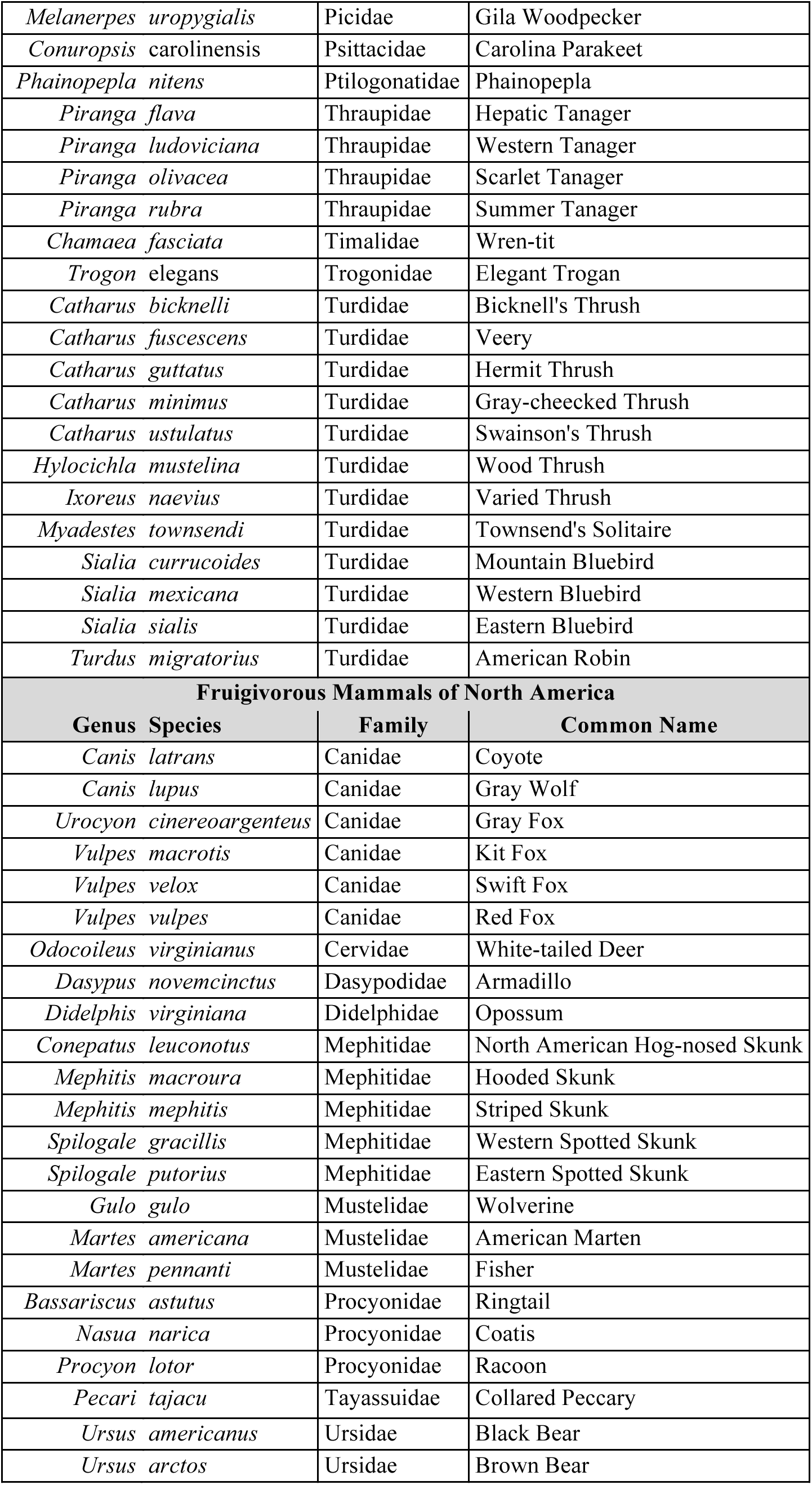
List of scatter-hoarding animals broken up into scatter-hoarding birds (top) and scatter-hoarding rodents (bottom). For both scatter-hoarding birds and mammals only one family possesses scatter-hoarding behavior (Corvidae and Rodentia respectively), therefore species are listed alphabetically by genus. Order of species does not insinuate a species ability to disperse seeds via scatter-hoar

**Table S4:**
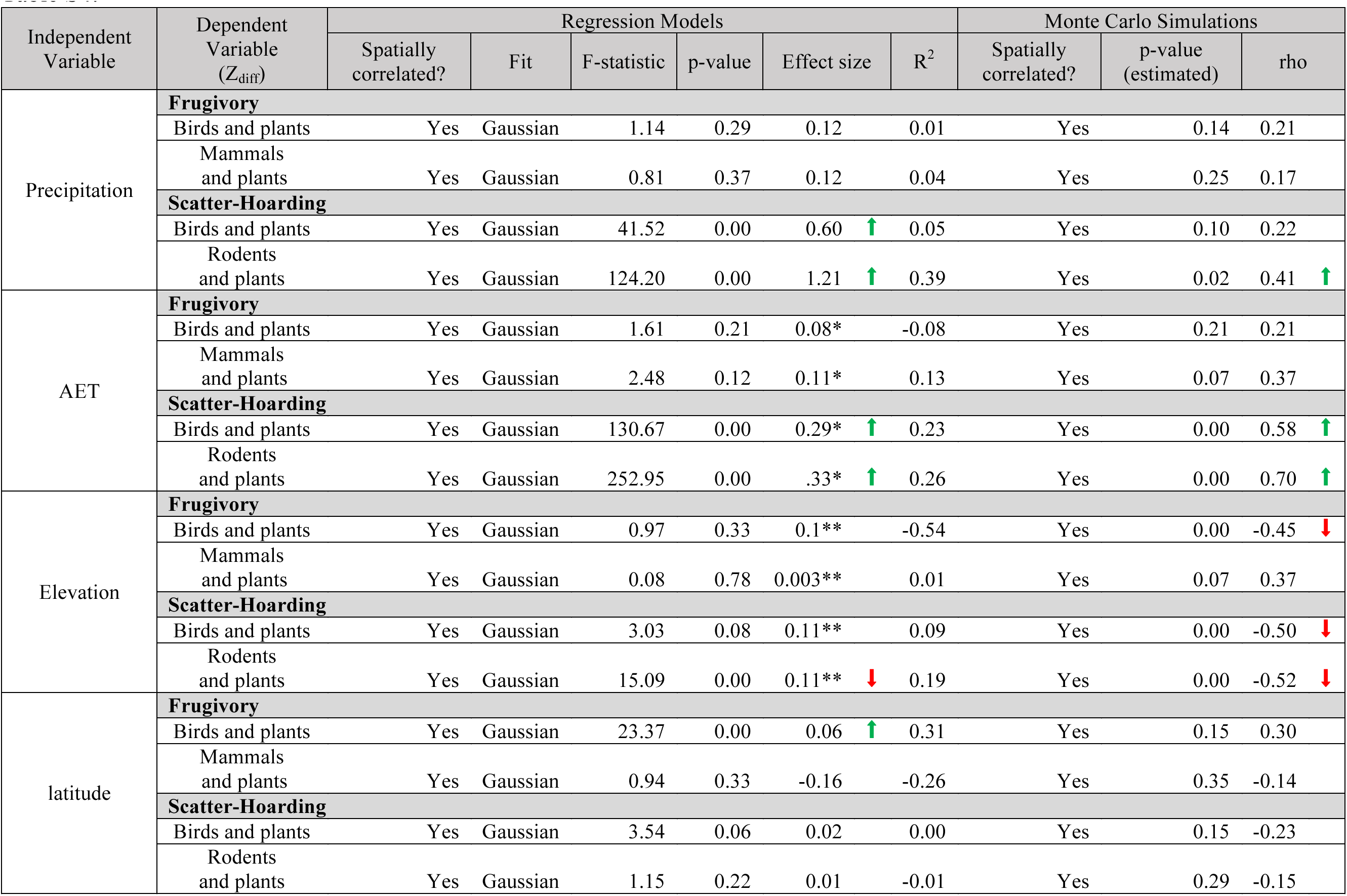
Statistical results of GLS models and Monte Caro simulations for mutualisms (Z_diff_) and environmental variables. An asterisk (*) signifies that the effect size is in units per 100 mm/yr AET and a double asterisk (**) signifies that the effect size is in units per 100 m elevation. Arrows signify if the relating ship is negative (downward arrow) or positive (upward arrow).

## SUPPLEMENTARY INFORMATION–SPATIAL RANDOMIZATIONS

I ncluded in this document are the results from the complete (S6–9) and structured (S10–13) spatial randomizations. The methods are described in *Materials and Methods, Data Analysis* of the paper, and the code to recreate the analysis and figures can be found at https://github.com/dispersing/SpatialRandomizations.

Figure S5 shows an example of an observed pattern (column 1) and 10 complete randomizations (rows 1 and 2) and 10 structured randomizations (rows 3 and 4). Figures S6–9 show the results for the complete randomizations of the correlations between animal and environmental variables (S5), the animal and plant variables (S7), the difference in richness and environmental variables (S8), and the plant and environmental variables (S9). Figures S10–13 show the results for the structured randomizations of the correlations between animal and environmental variables (S10), the animal and plant variables (S11), the difference in richness and environmental variables (S12), and the plant and environmental variables (S13).Variables names translate:

I. Environmental variables

a. aet: actual evapotranspiration
b. dem: elevation
c. ppt: precipitation
d. lat: latitude
II. Animal variables

a. a.mut: all mutualist animals
b. a.hoard: hoarding animal guild
c. a.frug: fruigivorous animal guild
d. a.hoard.rod: hoarding rodent subguild
e. a.hoard.bird: hoarding bird subguild
f. a.frug.mamm: frugivorous mammal subguild
g. a.frug.bird: frugivorous bird subguild
III. Plant variables

a. p.mut: all mutualist plants
b. p.hoard: hoarded plant guild
c. p.frug: fruigivory plant guild
d. p.hoard.rod: rodent-hoarded plant subguild
e. e.p.hoard.bird: bird-hoarded plant subguild
f. f.p.frug.mamm: mammal frugivory plant subguild
g. g.p.frug.bird: bird frugivory plant subguild
IV. Richness difference

a. d.mut: difference in richness of all mutualists
b. d.hoard: difference in richness of the hoarding guild
c. d.frug: difference in richness of the frugivory guild
d. d.hoard.rod: difference in richness of the rodent hoarding subguild
e. d.hoard.bird: difference in richness of the bird hoarding subguild
f. d.frug.mamm: difference in richness of the mammal frugivory subguild
g. d.frug.bird: difference in richness of the bird frugivory subguild

**Figure S5.**
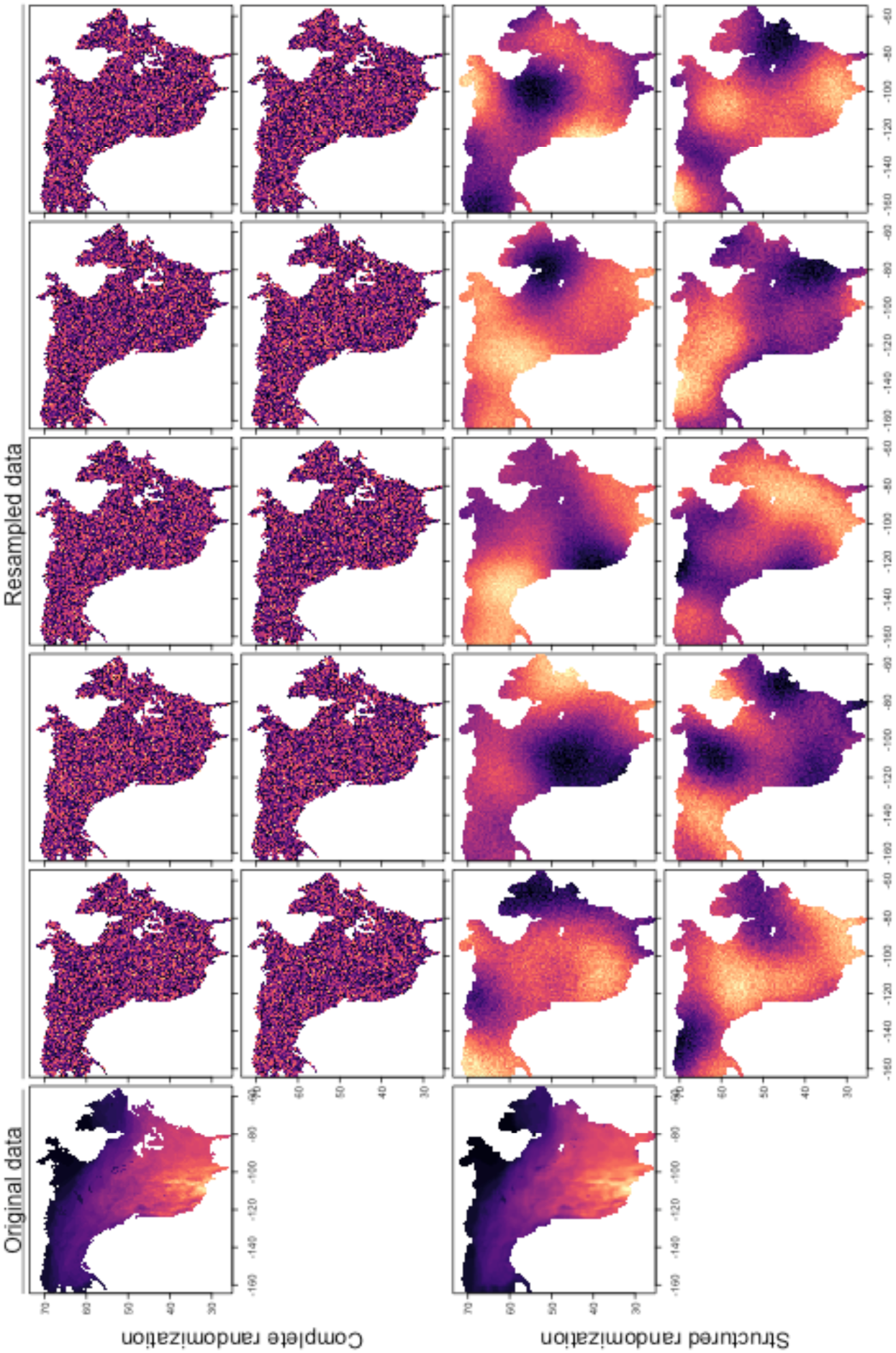
An example of an observed pattern (column 1, “Original data”) and 10 complete randomizations (rows 1 and 2) and 10 structured randomizations (rows 3 and 4). This example is to visually demonstrate to the reader the difference between the two tests. The latter preserves the spatial autocorrealtive structure and is therefore a more rigorous statistical test.

**Figure S6.**
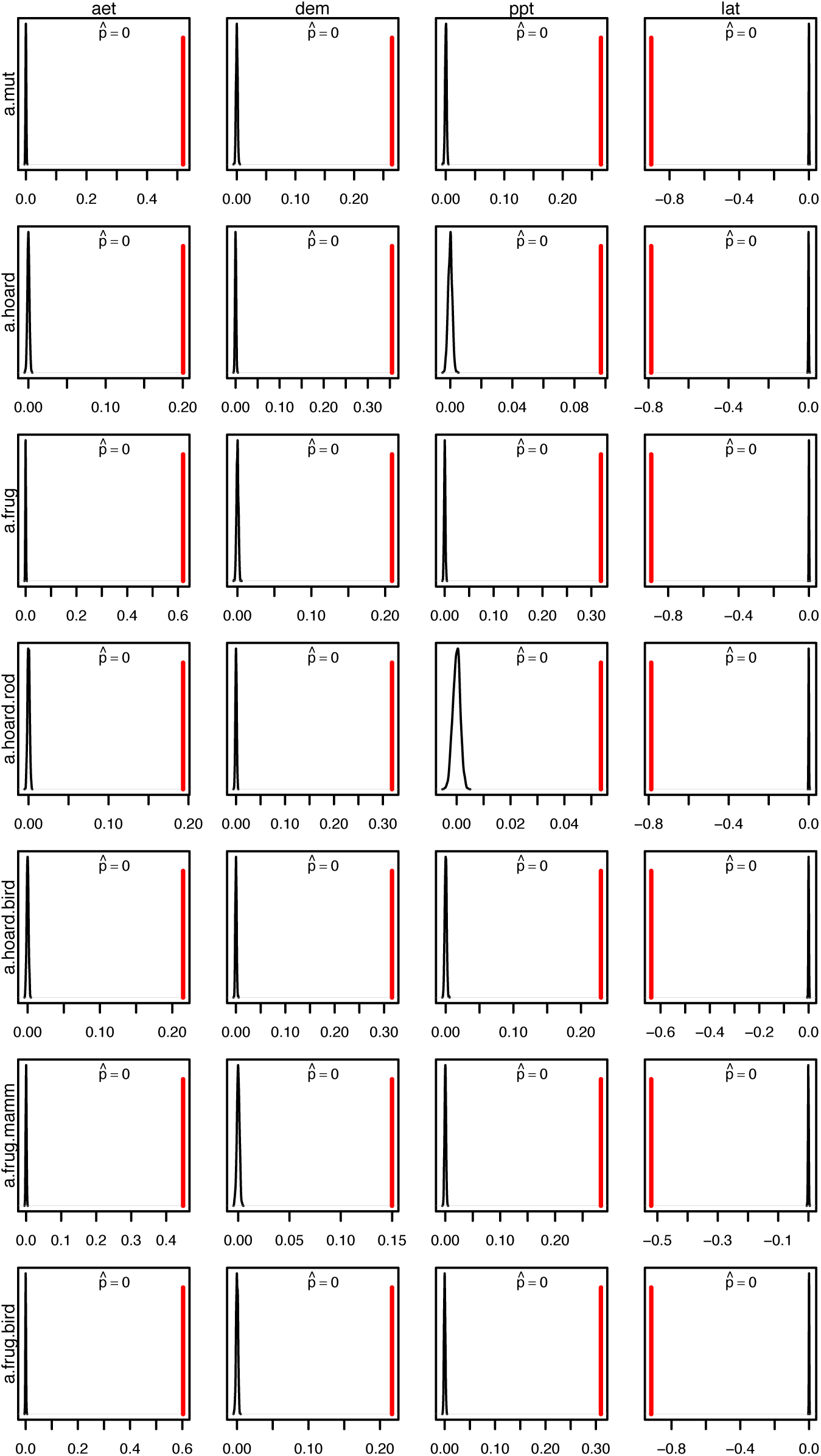
Results for the complete randomizations of the correlations between animal and environmental variables. Every randomization in each comparison had an observed value more extreme than all of randomizations, meaning all had an estimated *p*-value, 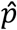=0.

**Figure S7.**
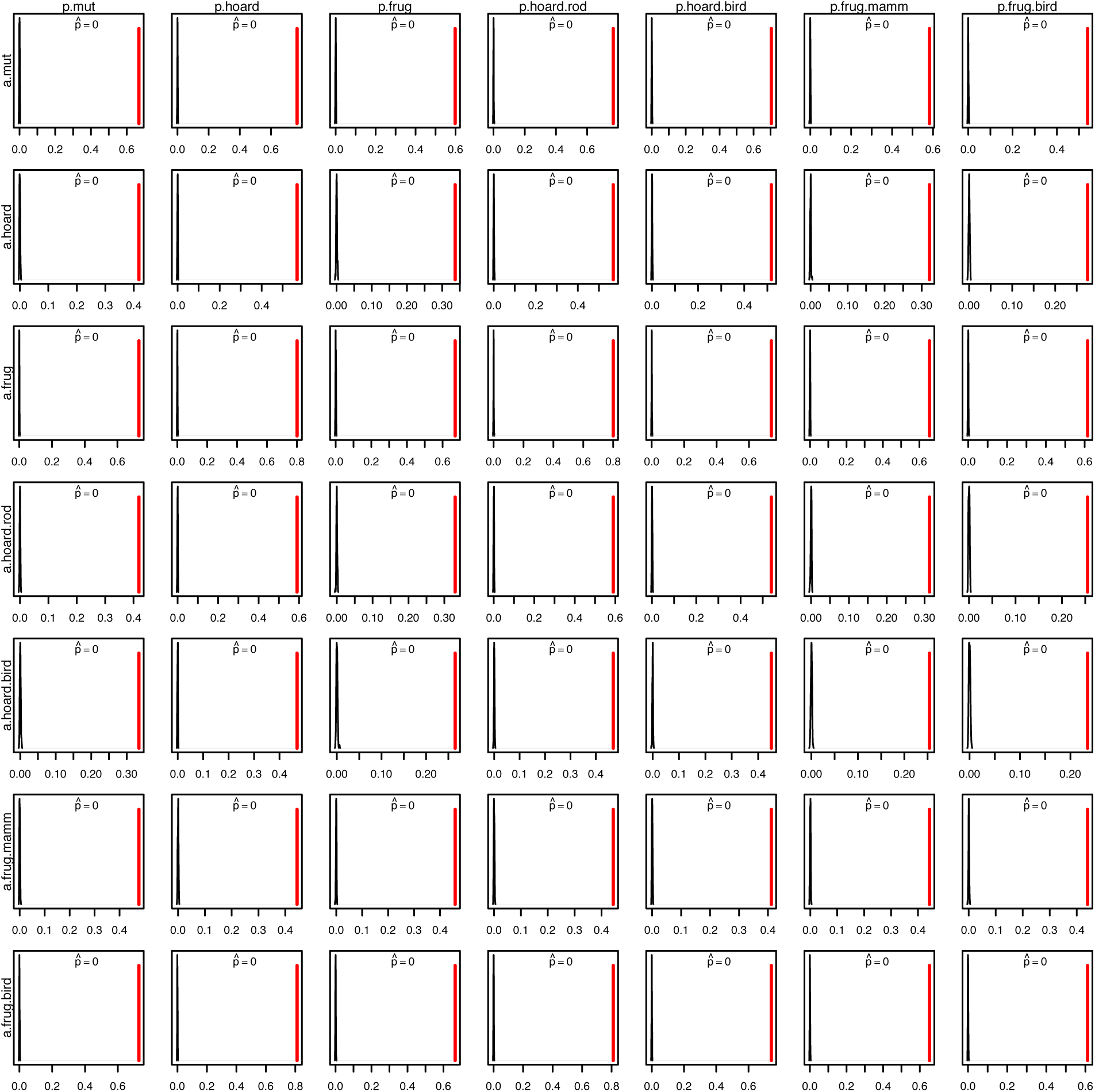
Results for the complete randomizations of the correlations between animal and plant variables. Every randomization in each comparison had an observed value more extreme than all of randomizations, meaning all had an estimated *p*-value, 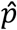=0.

**Figure S8.**
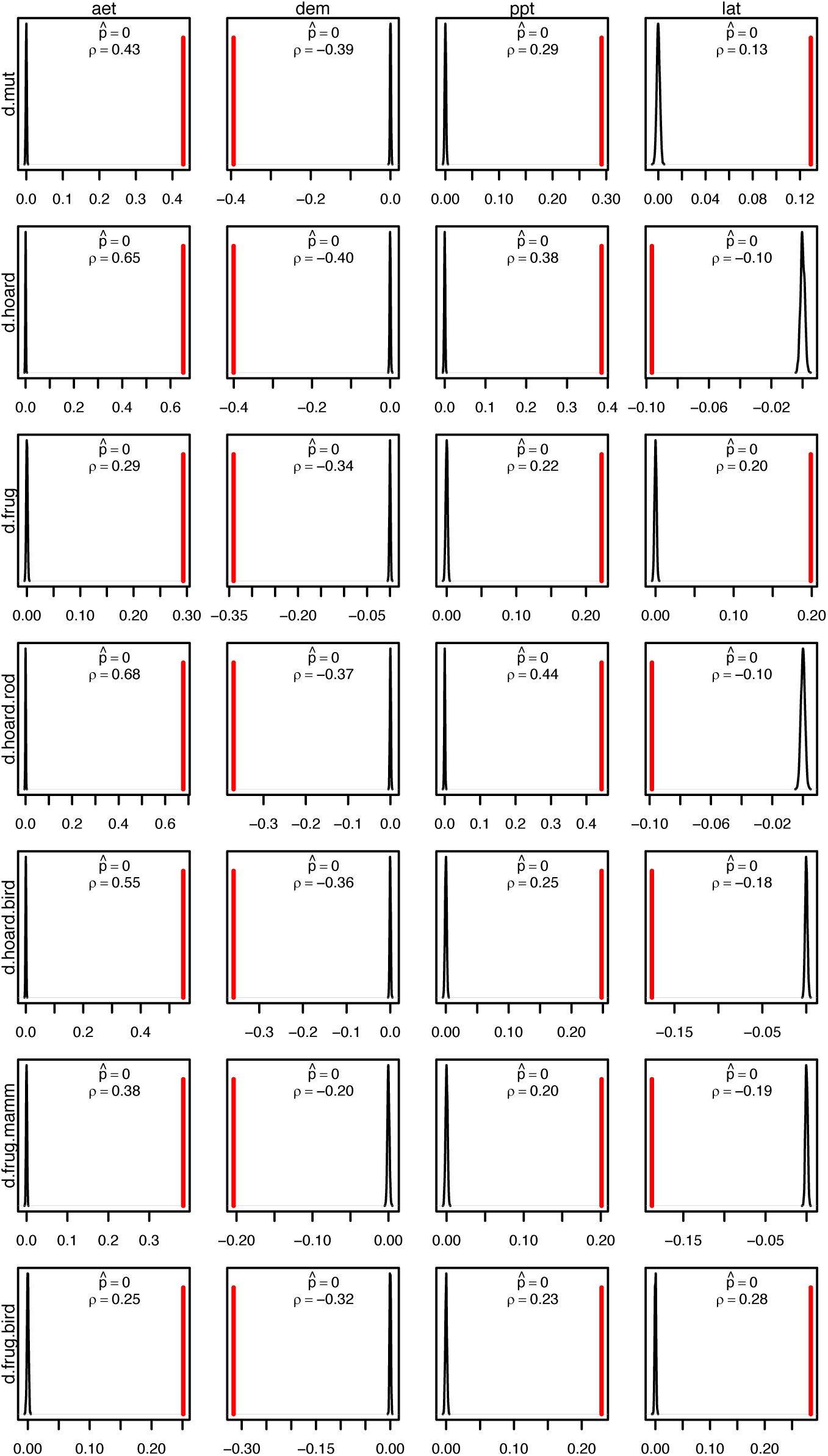
Results for the complete randomizations of the correlations between difference in richness and environmental variables. Every randomization in each comparison had an observed value more extreme than all of randomizations, meaning all had an estimated *p*-value, 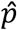=0.

**Figure S9.**
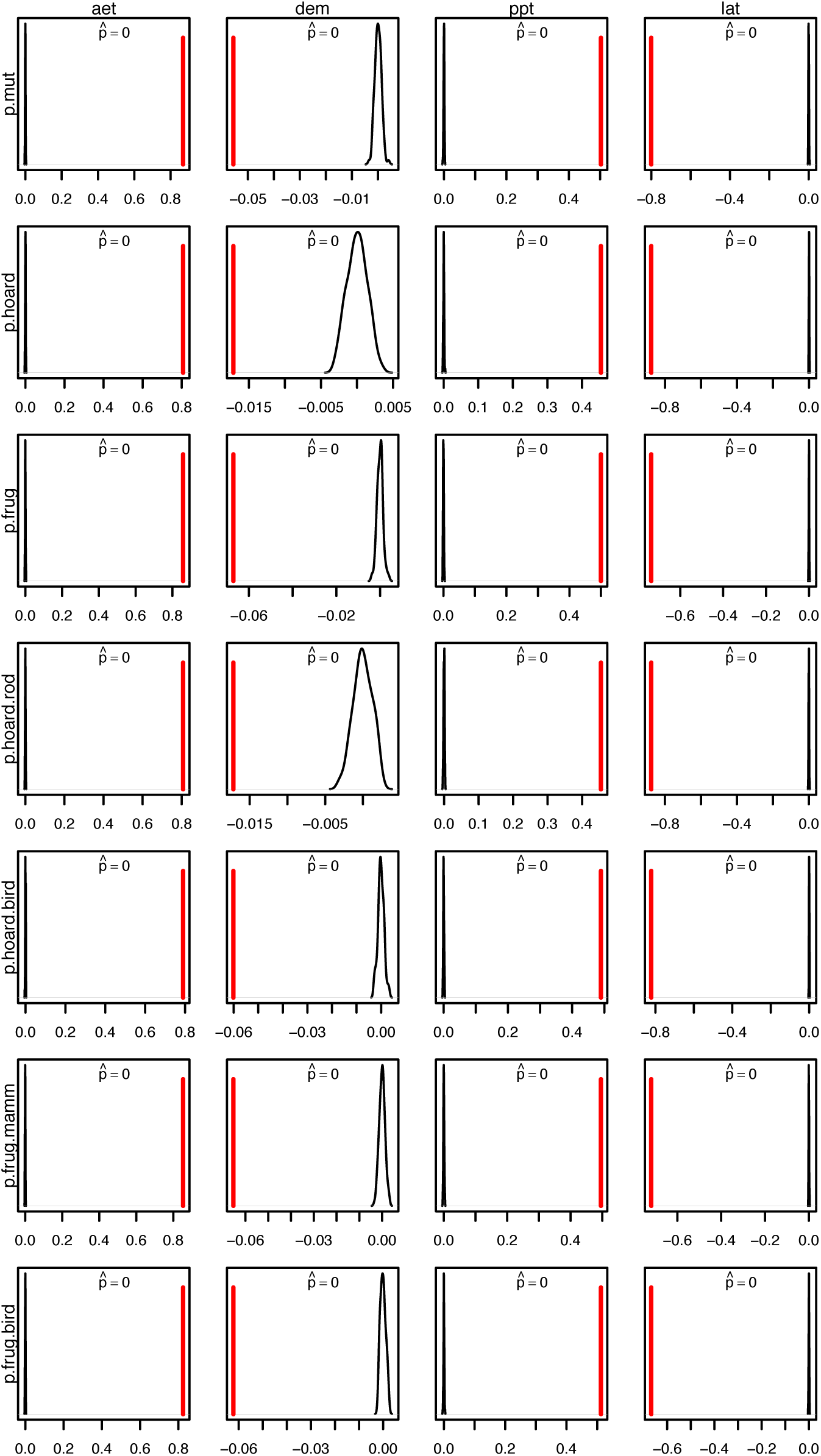
Results for the complete randomizations of the correlations between plant and environmental variables. Every randomization in each comparison had an observed value more extreme than all of randomizations, meaning all had an estimated *p*-value, 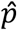=0.

**Figure S10.**
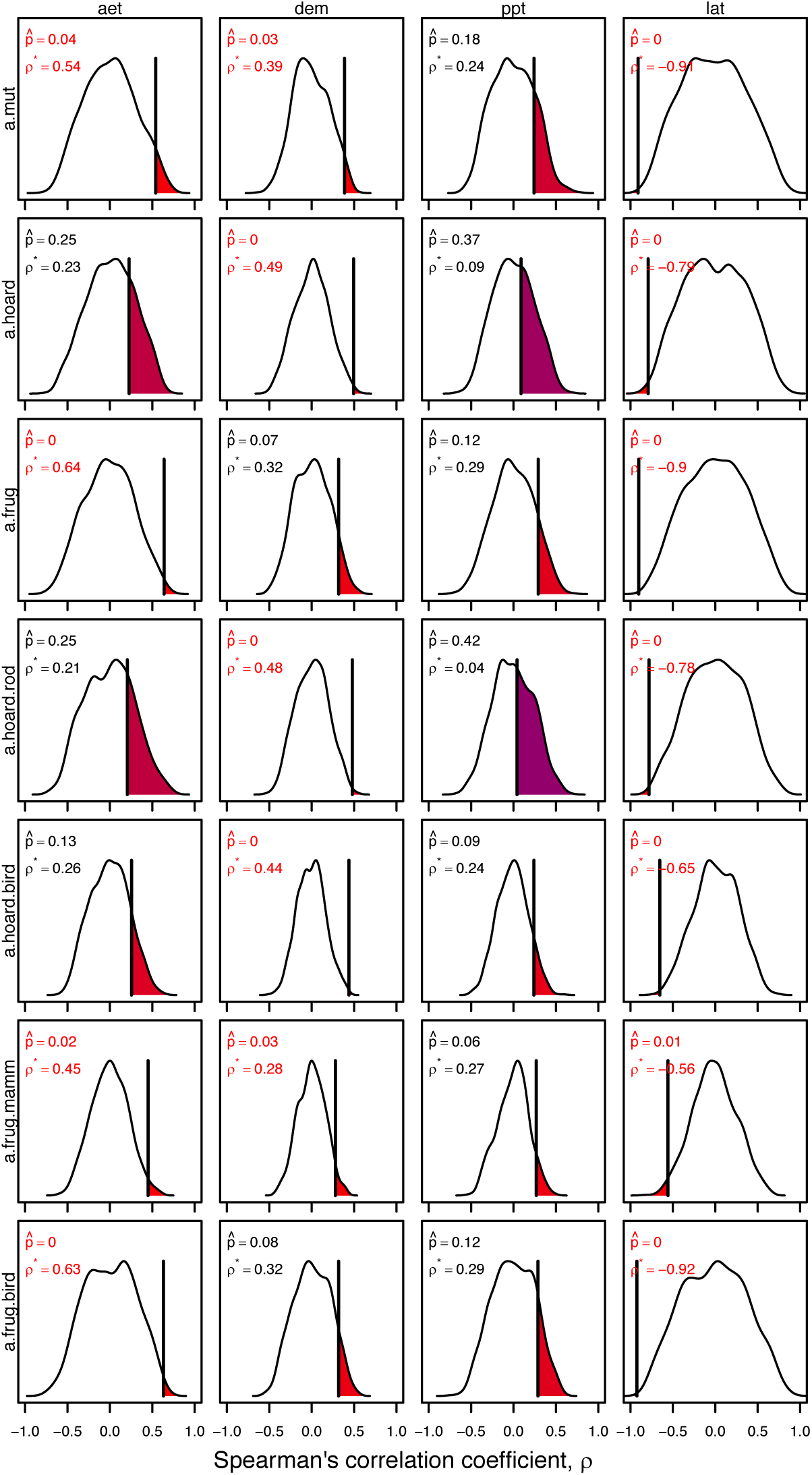
Results for the structured randomizations of the correlations between animal and environmental variables. Within each panel, the distribution represents the randomized Spearman’s correlation coefficient, *ρ*; vertical black lines represent the observed Spearman’s correlation coefficient, *ρ*^*^; shaded areas represent parts of the distribution that were more extreme than p*; the *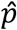* is the randomizations ≥ *ρ*^*^; and if ≤ 5% of the randomizations were more extreme than *ρ*^*^ then we interpret that as statistically significant and color the statistical font red rather than black.

**Figure S11.**
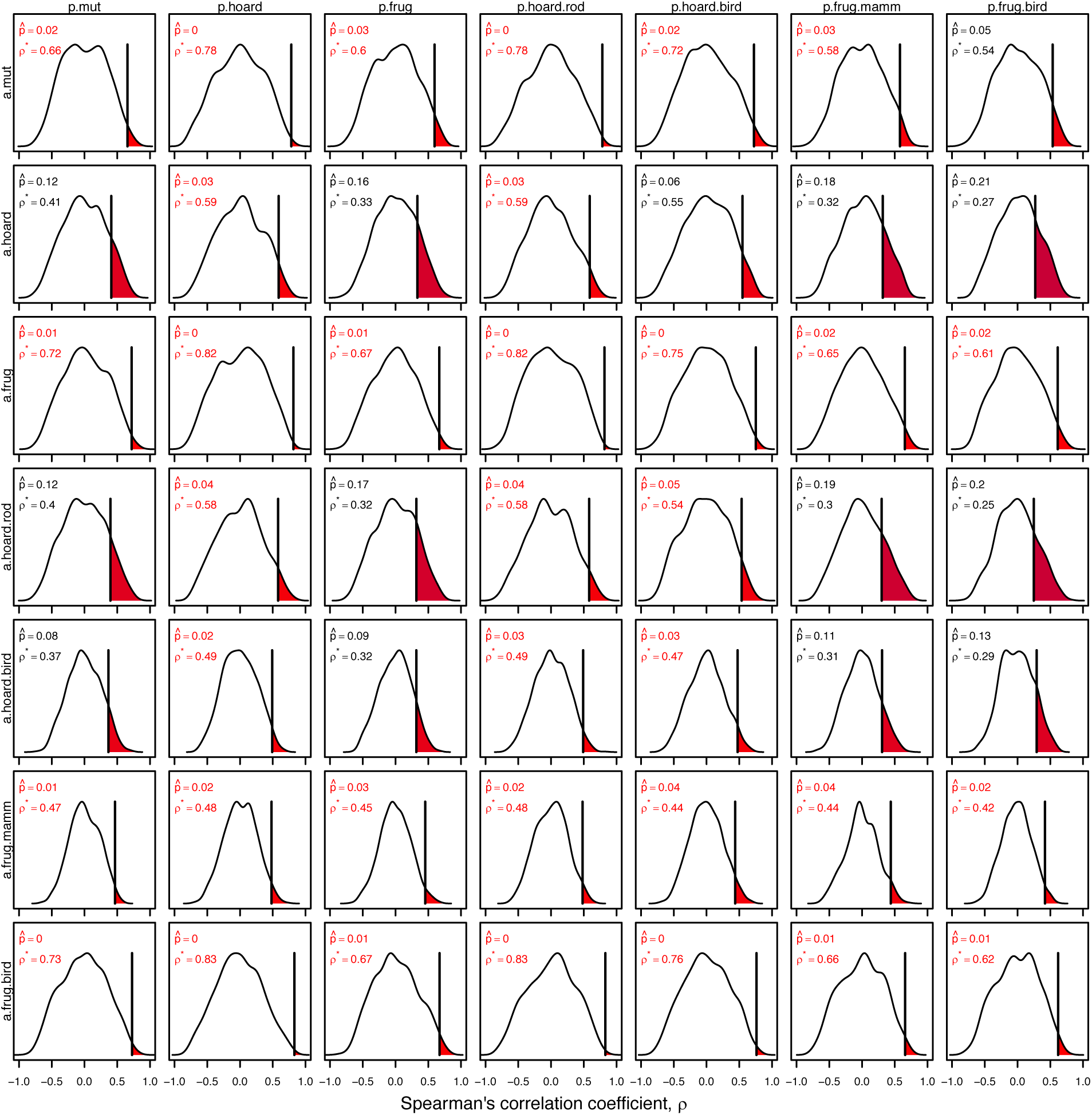
Results for the structured randomizations of the correlations between animal and plant variables. Within each panel, the distribution represents the randomized Spearman’s correlation coefficient, ρ; vertical black lines represent the observed Spearman’s correlation coefficient, *ρ^*^;* shaded areas represent parts of the distribution that were more extreme than *ρ*;* the *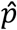* is the randomizations ≥ *ρ*^*^; and if ≤ 5% of the randomizations were more extreme than *ρ*^*^ then we interpret that as statistically significant and color the statistical font red rather than black.

**Figure S12.**
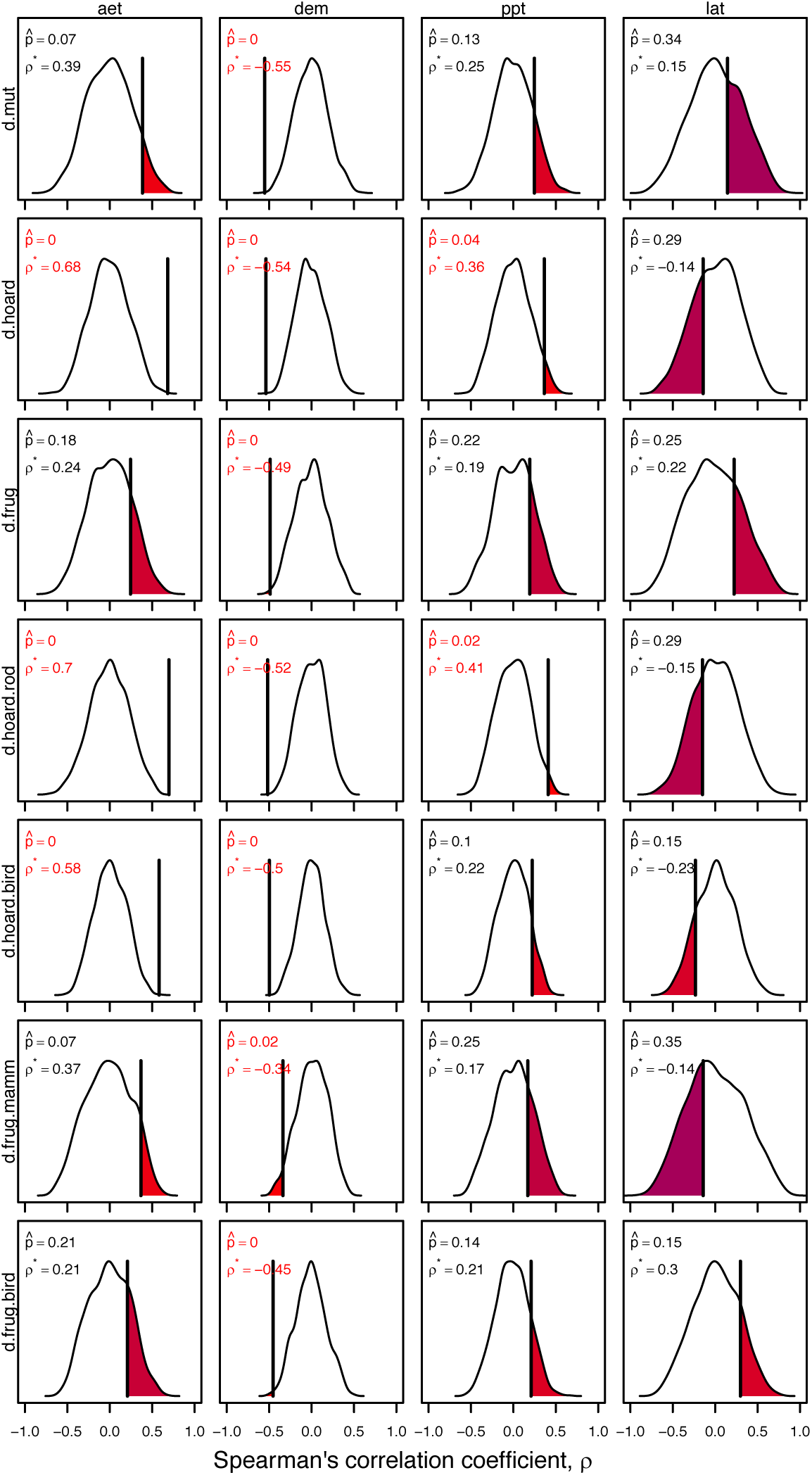
Results for the structured randomizations of the correlations between the difference in richness and environmental variables. Within each panel, the distribution represents the randomized Spearman’s correlation coefficient, *ρ*; vertical black lines represent the observed Spearman’s correlation coefficient, *ρ*^*^; shaded areas represent parts of the distribution that were more extreme than *ρ*^*^; the *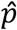* is the randomizations ≥ *ρ*^*^; and if ≤ 5% of the randomizations were more extreme than *ρ*^*^ then we interpret that as statistically significant and color the statistical font red rather than black.

**Figure S13.**
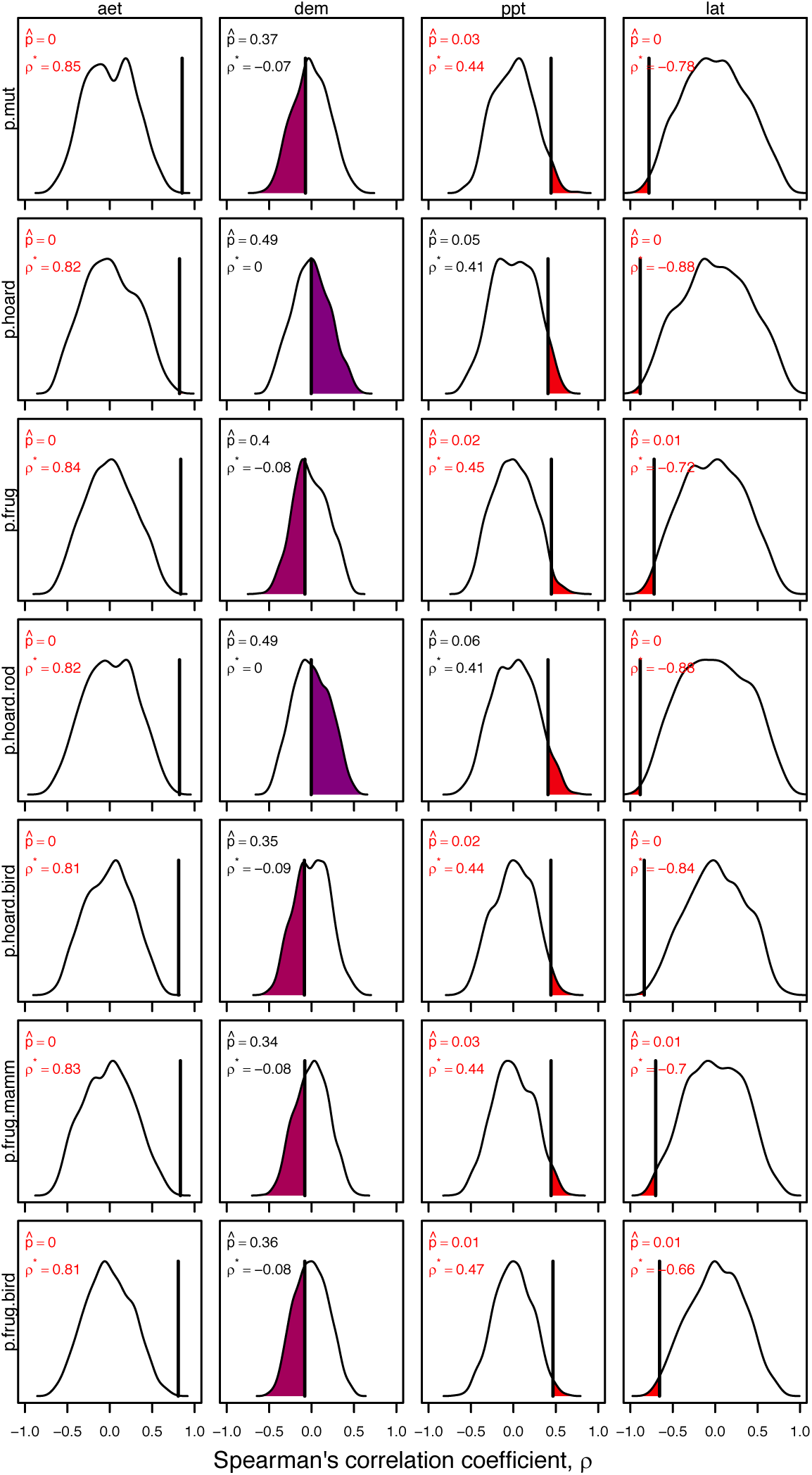
Results for the structured randomizations of the correlations between the plant and environmental variables. Within each panel, the distribution represents the randomized Spearman’s correlation coefficient, *ρ*; vertical black lines represent the observed Spearman’s correlation coefficient, *ρ*^*^; shaded areas represent parts of the distribution that were more extreme than *ρ*^*^; the *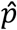* is the randomizations ≤ *ρ*^*^; and if ≤ 5% of the randomizations were more extreme than *ρ*^*^ then we interpret that as statistically significant and color the statistical font red rather than black.

